# Mating proximity blinds threat perception

**DOI:** 10.1101/2024.04.23.590677

**Authors:** Laurie Cazalé-Debat, Lisa Scheunemann, Megan Day, Tania Fernandez-d.V. Alquicira, Anna Dimtsi, Youchong Zhang, Lauren A Blackburn, Charles Ballardini, Katie Greenin-Whitehead, Eric Reynolds, Andrew C Lin, David Owald, Carolina Rezaval

## Abstract

Romantic engagement can bias sensory perception. This ’love blindness’ reflects a common behavioral principle across organisms: favoring pursuit of a coveted reward over potential risks. In the case of animal courtship, such sensory biases may support reproductive success but can also expose individuals to danger, such as predation. How do neural networks balance the trade-off between risk and reward? Here, we discover a dopamine-governed filter mechanism in male *Drosophila* that reduces threat perception as courtship progresses. We show that during early courtship stages, threat-activated visual neurons inhibit central courtship nodes via specific serotonergic neurons. This serotonergic inhibition prompts flies to abort courtship when they see imminent danger. However, as flies advance in the courtship process, the dopaminergic filter system reduces visual threat responses, shifting the balance from survival to mating. By recording neural activity from males as they approach mating, we demonstrate that progress in courtship is registered as dopaminergic activity levels ramping up. This dopamine signaling inhibits the visual threat detection pathway via Dop2R receptors, allowing male flies to focus on courtship when they are close to copulation. Thus, dopamine signaling biases sensory perception based on perceived goal proximity, in order to prioritize between competing behaviors.

## MAIN

Every day, animals make decisions that require balancing opportunities and risks. Research has explored this arbitration in hypothetical scenarios^1,2^, as well as in conflicting behavioral and decision-making contexts in humans^3^, rodents^4–6^, and invertebrates^7–13^. However, we still lack a detailed mechanistic understanding of how conflict is resolved in the brain, particularly when the dangers and benefits are crucial life choices.

Most organisms face a perpetual trade-off between survival and reproduction. While courting a mate increases the chances of reproductive success, it also carries risks, such as exposure to predation. Recent work has revealed how sex drive and threat avoidance are independently signaled in the brain^14–22^,yet it remains unclear how these needs are prioritized when they are in conflict. Avoiding threats can be a life-saving decision, but excessive caution might result in missed mating opportunities. This raises the question of how animals suppress courtship when it is better to run away, and how this is reversed when the rewards of courtship outweigh the risk of predation (e.g., if mating is imminent).

The role of dopaminergic signaling as a keystone in the architecture of motivation, need, and reward is well-established across species^23–26^. Beyond its classical reward-related functions, research suggests that dopamine plays a key role in relaying the value of sensory input and internal/behavioral states to decision-making centers in the brain, thus adapting behavior^12,27–41^. Yet, how dopamine dynamically modulates sensory valence and influences decision-making during conflict remains poorly understood. A potential mechanism may be offered by sensory filter systems^42–44^ which eliminate superfluous inputs and highlight relevant information to facilitate appropriate behaviors. Such filtering systems could thus serve as a means to shut down competing sensory inputs when animals are close to achieving a crucial goal. Here, we describe a state-dependent filter system driven by dopamine that allows *Drosophila* males to filter out external threats and focus on courtship when they are close to mating.

### Threat-driven visual input disrupts early courtship via LC16 visual neurons

*Drosophila* males engage in a series of stereotyped, progressive courtship steps that include tapping the female with their legs, vibrating a wing to produce a species-specific song, contacting her genitalia with their mouthparts, and bending the abdomen to attempt copulation^18,45,46^ (Fig. 1a, Supplementary Videos 1-2). If the female is receptive, the male typically exhibits a strong motivation and commitment to courtship^17,45^ persisting until copulation is achieved. Yet, what happens when the urge to court is fraught with risk?

**Figure 1:**
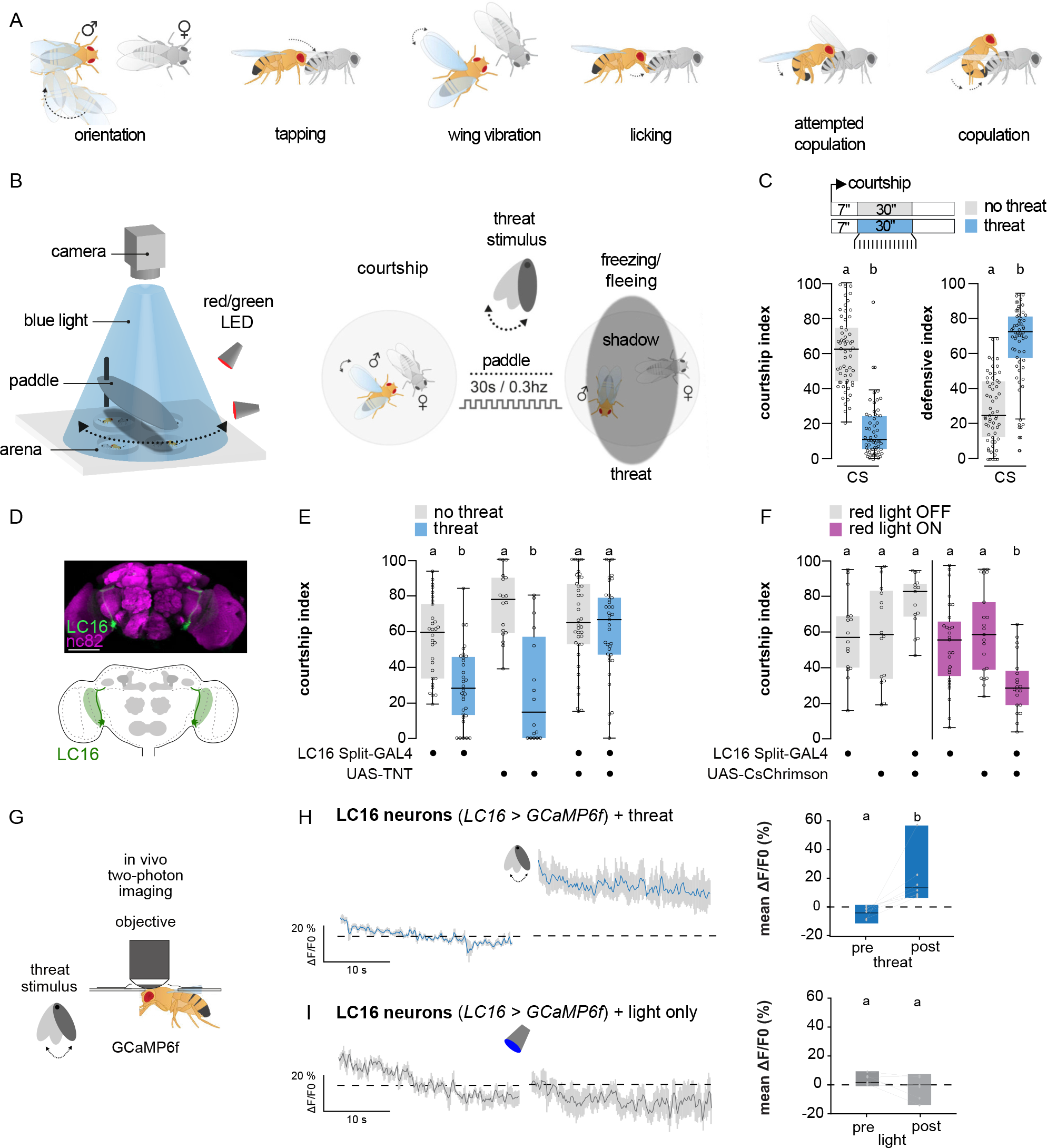
Courtship is disrupted by visual predatory threats in male flies via LC16 neurons. **(A)** Schematic of the *Drosophila* courtship ritual. **(B)** Schematic of the action selection paradigm: A naive male courting an immobile virgin female for at least 7 s is exposed to a visual threat created by repeatedly passing an object which casts an overhead shadow at 0.3 Hz for 30 s. In response to the visual threat, males interrupt courtship and display defensive behaviors (e.g. freezing and escaping). **(C)** Courtship and defensive behaviors of 5-7 day old wild-type Canton S (CS) males in the absence or presence of the visual threat (n=59 for both). Control and experimental flies were both assessed 7 s after they started courting (Fig. 1c upper panel). Courtship and defensive indexes were calculated as time the male spent displaying a given behavior x 100 / duration of the threat delivery (30 s). **(D)** Upper panel: UAS-mCD8-GFP (green) expressed under the control of LC16 Split- GAL4 in the adult brain; neuropil counterstaining with anti-nC82 (purple). Scale bar: 50 μm. Bottom panel: Schematic of LC16 lobula columnar (LC) neurons morphology. Somata are located near the ventral tip of the lobula of the optic lobe and project their dendrites and axons within the lobula neuropil and the PVLP glomeruli within the central brain. **(E)** Courtship index of males in which LC16 neurons have been silenced with tetanus toxin light chain (TNT), as well as genetic controls, in the absence (grey plots, n=18-38) or presence of the threat (blue plots, n=16-37). **(F)** Courtship index after artificial activation of LC16 neurons under 660 nm light using CsChrimson in the absence of the threat (n^red^ ^light^ ^OFF^=15-16; n^red^ ^light^ ^ON^=20-27) and genetic and light controls. **(G)** Schematic of the two-photon functional imaging set-up with threat delivery. The male is tethered to an imaging chamber that allows access of the objective to the brain but leaving the eyes functional. A motorized paddle and light source are installed underneath the imaging chamber, directing a movable shadow in front of the head of the fly keeping the same parameters used during the behavioral threat assay (0.3 Hz for 30 s). **(H)** Fluorescence intensity trace (ΔF/F0) over time of male flies expressing GCaMP6f in LC16 neurons pre and post 30 s of threat exposure under the microscope. Right: Quantification of mean ΔF/F0 comparing the 30 s pre and post time windows (n=7). **(I)** Fluorescence intensity trace (ΔF/F0) over time of male flies expressing GCaMP6f in LC16 neurons pre and post 30 s of light exposure without threat under the microscope. Right: Quantification of mean ΔF/F0 comparing the 30 s pre and post time windows (n=6).

To dissect the neural circuitry that prioritizes between these competing needs, we first established a sex-danger conflict model, where courting *Drosophila* males were presented with a visual threat: a predator-like moving shadow^47^. Indeed, in the absence of females, the threat caused males to show defensive responses such as running away and freezing^47^ (Fig. 1b; Extended Data Fig. 1a, b). To eliminate confounding effects of female behavior, we used immobile virgin females. As expected, in the absence of the threat, wild-type males vigorously courted the females and showed low defensive behaviors (courtship index (CI)=62%; defensive index (DI)=24%; Fig. 1c, Extended Data Fig. 1c, Supplementary Video 1). However, upon threat presentation, males immediately halted the courtship and engaged in defensive responses (CI=20%, DI=72%, Fig. 1c, Extended Data Fig. 1c, Supplementary Videos 3-4).

We next sought to determine which neurons detect the visual threat signal. Lobular columnar (LC) visual projection neurons connect early visual processing with central brain areas and respond to ethologically relevant visual features, such as conspecifics and motion signals^16,48–50^. We therefore hypothesized that LC neurons might detect and convey visual threats to higher brain centers to inhibit courtship. Consistent with this, we found that LC16 neurons^51–53^ were required to prioritize defensive responses over courtship (Fig. 1d,e, Extended Data Fig. 1d,e). When LC16 neurons were silenced by expressing the tetanus toxin light chain (TNT), males courted the female despite the visual threat (CI^nothreat^=64% vs. CI^threat^=66%, Fig. 1e, Extended Data Fig. 1d). Conversely, when LC16 neurons were optogenetically activated using the red channelrhodopsin CsChrimson in the absence of the threat, males stopped courting, mimicking the effect of the visual threat (CI^lightOFF^=84% vs. CI^lightON^=30%, Fig. 1f, Extended Data Fig.1e). Blocking LC16 synaptic output disrupted visual threat responses in solitary males (Extended Data Fig.1f) but did not alter responses to mechanical threats in courting males (Extended Data Fig. 1g). This confirms that LC16 neurons are implicated in defensive behaviors in response to threatening visual features, agreeing with previous results^52,53^. Males with silenced LC16 neurons showed normal courtship behaviors in the absence of threats (CI^nothreat^=64% vs. CI^threat^=66%, Fig. 1e grey plot). Therefore, LC16 neurons suppress courtship in response to visual threats by modulating courtship-related circuits, rather than directly controlling these behaviors.

LC16 neurons have been shown to be sensitive to both looms and moving bars^52–54^. To verify that LC16 neurons can also perceive the moving shadow stimulus, we performed *in vivo* two-photon calcium imaging (Fig. 1g). Threat exposure induced a substantial calcium influx in LC16 neurons (+15% in ΔF/F0, Fig. 1h). In contrast, non-threat visual stimuli, such as light-only controls or female sensory cues, elicited no detectable response (ΔF/F0=0%, Fig. 1i, Extended Data Fig. 1h, Extended Table 2). Altogether, our findings show that LC16 neurons detect visual threats and prompt the flies to cease courtship and engage in defensive actions.

### 5-HT neurons are required to prioritize survival over courtship

Previous research in fish and mammals has shown that serotonin (5-HT) not only modulates fight-or-flight responses associated with predators but also increases due to factors such as social stress^55,56^. Therefore, we postulated that inhibition of courtship could be driven by 5-HT signaling. To test this hypothesis, we either blocked all 5-HT neurons or blocked 5-HT synthesis altogether using RNAi against tryptophan hydroxylase (TRH), the enzyme involved in serotonin synthesis. In both forms of inhibition, males—whether solitary or paired with a female—did not increase their speed or cease courtship in the presence of the threat (Extended Data Fig. 2a-f). In contrast, when the 5-HT neurons were optogenetically activated, courtship halted in the absence of the threat (Extended Data Fig. 2g,h). We conclude from these experiments that 5-HT neurons are important for prioritizing escape responses over courtship in response to visual threats.

### Visually-driven 5-HT signaling inhibits central courtship nodes

In our search for the neurons involved in the courtship/escape choice, we examined the P1 cluster, a central mating-regulation hub that initiates courtship in response to female sensory cues and internal states^16,18,22,57–65^ (Fig. 2a). Given that P1 neurons integrate other competing drives^8,64,66,67^ (e.g., aggression, sleep and feeding), we hypothesized that they may also be implicated in the courtship/escape choice. Indeed, optogenetically activating P1 neurons in males during the exposure to the visual threat caused them to continue to court, overriding the threat response (CI^threat+redlightON^=44% for P1>Chrimson vs. CI^threat+redlightON^=8-20% for genetic control flies, Fig. 2b, Extended Data Fig. 3a). This result suggests that visual threats might block courtship by inhibiting P1 neurons.

**Figure 2:**
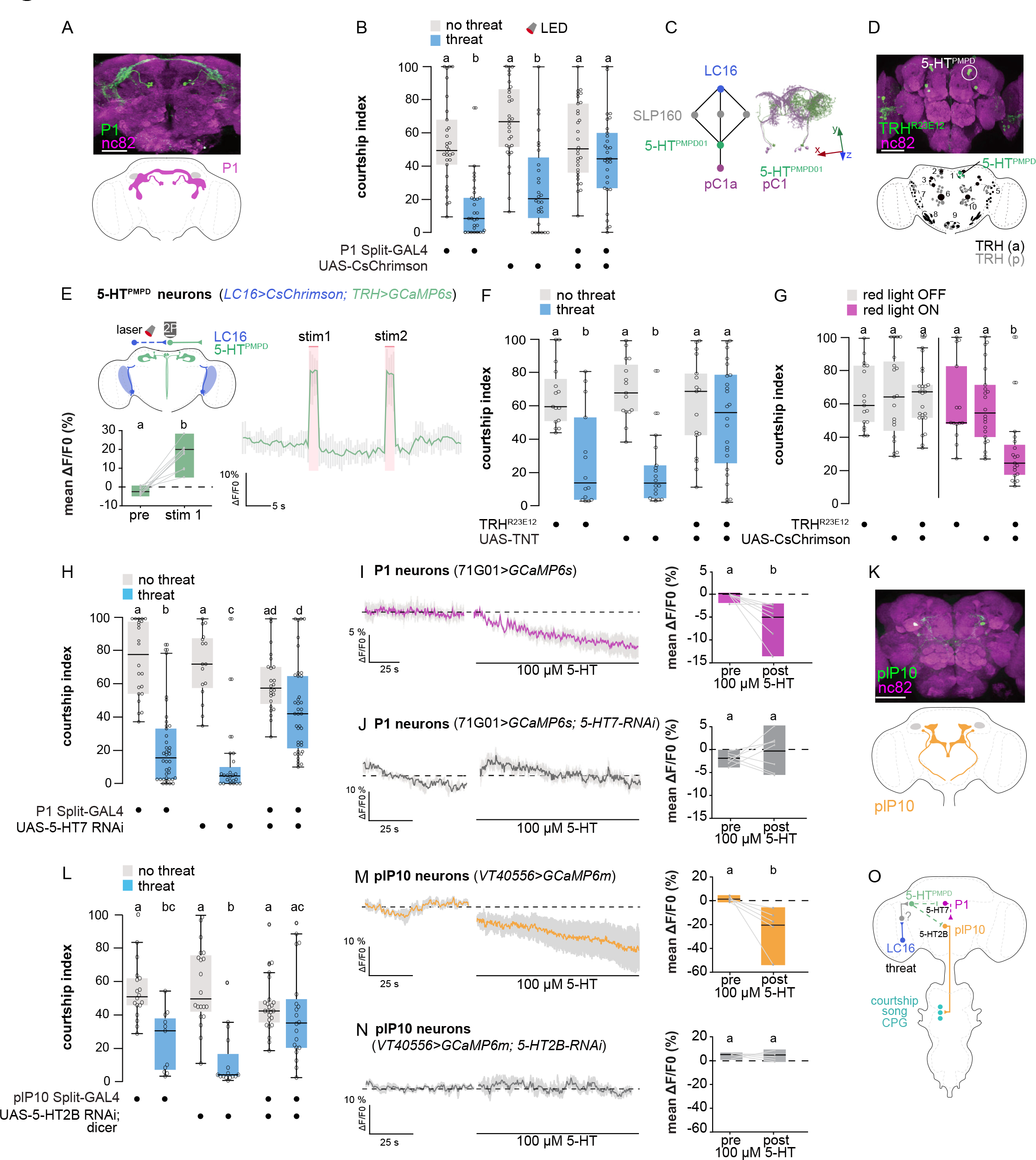
Visually-driven 5-HT signalling inhibits P1 and plP10 courtship-promoting neurons. **(A)** Upper panel: UAS-mCD8-GFP (green) expressed under the control of P1 Split-GAL4 in the adult brain; anti-nC82 (purple). Scale bar: 50 μm. Bottom panel: Schematic of the P1 courtship cluster in the brain. **(B)** Courtship index after artificial activation of P1 neurons under 660 nm red light using CsChrimson in absence (grey plots, n=30) or presence (blue plots, n=28-30) of the threat. **(C)** Left panel: neural path connecting LC16 VPNs to female pC1a neuron through 5- HT^PMPD01^ neuron predicted by the hemibrain connectome^68^. Right panel: electron microscope reconstruction of images of pC1(a,b,e) neurons and 5-HT^PMPD01^ neurons in the female central nervous system. x,y,z arrows represent the anterior/posterior, medial/lateral and dorsal/ventral axis, respectively. **(D)** Upper panel: Genetic intersection using the Split-GAL4 system (R23E12-AD; TRH- DBD) to target the PMPD cluster. Bottom panel: schematic representation^69^ of the different anterior (black, ‘a’) and posterior (grey, ‘p’) serotonin clusters in the adult central brain. 1: PMPD (posterior medial dorsal protocerebrum), 2: ADMP (anterior dorsomedial protocerebrum), 3: ALP (anterior lateral protocerebrum), 4: PMPM (posterior medial protocerebrum medial), 5: PLP (posterior lateral protocerebrum), 6: AMP (anterior medial protocerebrum), 7: LP (lateral protocerebrum), 8: SEL (lateral subesophageal ganglion), 9: SEM (medial subesophageal ganglion), 10: PMPV (posterior medial protocerebrum ventral). **(E)** Left panel: schematic of LC16 visual projection neurons expressing CsChrimson and TRH^PMPD^ neurons expressing the calcium sensor GCaMP6s. Right panel: Fluorescence intensity trace (ΔF/F0) over time of males expressing GCaMP6s in 5HT^PMPD^ neurons after artificial activation of LC16 visual neurons using red-shifted channelrhodopsin CsChrimson under 1040 nm two-photon laser stimulation. Lower panel: Quantification of mean ΔF/F0 of baseline activity compared to first stimulation window (n=9). **(F)** Courtship index of males while silencing TRH^R23E12^ neurons using TNT as well as genetic controls in the absence (grey plots, n=15-17) or presence of the threat (blue plots, n=14-24). **(G)** Courtship index after artificial activation of TRH^R23E12^ neurons under 660 nm light using CsChrimson in absence of the threat (n^red^ ^light^ ^OFF^=17-25; n^red^ ^light^ ^ON^=14-19) and their genetic and light controls. **(H)** Courtship index of males with the 5-HT7 receptor knocked down specifically in P1 neurons using small RNA interference, and genetic controls in absence (grey plots, n=16-24) or presence of the threat (blue plots, n=24-39). **(I)** Fluorescence intensity trace (ΔF/F0) over time of male flies expressing GCaMP6s in P1 neurons pre and post application of 100 µM 5-HT. Right: Quantification of mean ΔF/F0 comparing the 30 s pre and last 30 s post time windows (n=7). **(J)** Fluorescence intensity trace (ΔF/F0) over time of male flies expressing GCaMP6s in P1 neurons with simultaneous knock down of the 5-HT7 receptor pre and post application of 100 µM 5-HT. Right: Quantification of mean ΔF/F0 comparing the 30 s pre and last 30 s post time windows (n=7). **(K)** Upper panel: UAS-mCD8-GFP (green) expressed under the control of plP10Split- GAL4 in the adult brain; neuropil counterstained with anti-nC82 (purple). Scale bar: 50 μm. Bottom panel: Schematic of plP10 neurons morphology within the central brain (1 single neuron per hemibrain). plP10 soma is located within the medial posterior brain innervating the lateral protocerebral complex and sends axon down to the anterior wing neuropil of the mesothoracic ganglion. **(L)** Courtship index of males with the 5-HT2B receptor knocked down specifically in plP10 neurons using RNA interference, and genetic controls, in absence (grey plots, n=18-24) or presence of the threat (blue plots, n=11-20). **(M)** Fluorescence intensity trace (ΔF/F0) over time of male flies expressing GCaMP6m in plP10 neurons pre and post application of 100 µM 5-HT. Right: Quantification of mean ΔF/F0 comparing the 30 s pre and last 30 s post time windows (n=6). **(N)** Fluorescence intensity trace (ΔF/F0) over time of male flies expressing GCaMP6m in plP10 neurons and simultaneous knock down of the 5-HT2B receptor pre and post application of 100 µM 5-HT. Right: Quantification of mean ΔF/F0 comparing the 30 s pre and last 30 s post time windows (n=6). **(O)** Network model underlying the escape and courtship choice. Dashed lines indicate indirect connection. Solid lines indicate direct connection. CPG: central pattern generator.

Using the female *Drosophila* connectome^68^, we identified a serotonergic (5-HT) neuron (5- HT^PMPD01^) in the posterior medial dorsal (PMPD) cluster (Fig. 2c, d) that receives input from LC16 neurons through an intermediate neuron and that in turn forms presynaptic connections on pC1, the female equivalent of P1 (Fig. 2c). Although it is not guaranteed that the same connection exists on male P1 neurons, thanks to the sexual dimorphism of pC1/P1, this connectome data made 5-HT^PMPD^ neurons attractive candidates to carry threat signals from LC16 to P1.

To assess whether LC16 and 5-HT^PMPD^ neurons are functionally connected, we optogenetically activated LC16 with CsChrimson and monitored 5-HT^PMPD^ responses by GCaMP6s calcium imaging (Fig. 2e). LC16 stimulation reliably increased the GCaMP6s signal in 5-HT^PMPD^ neurons (+22% in ΔF/F, Fig. 2e), while light stimulation in flies lacking CsChrimson had a slight, but not significant effect (Extended Data Fig. 3b, Extended Table 2), placing the 5- HT^PMPD^ cluster downstream of LC16 threat neurons. Furthermore, threat exposure triggered significant calcium influx in 5-HT^PMPD^ neurons (Fig. 5k), similar to the threat response of LC16 neurons.

To test the role of 5-HT^PMPD^ neurons in threat-induced suppression of courtship, we used a Split-GAL4 (TRH^R23E12^) targeting the 5-HT^PMPD^ cluster and a subset of 5-HT^LP^ neurons^69^ (Fig. 2d). Blocking TRH^R23E12^ synaptic output prevented solitary males from escaping threats (Extended Data Fig. 3c), implicating them in defensive responses. We further found that silencing TRH^R23E12^ neurons by either expressing TNT or knocking down the TRH enzyme led to persistent courtship activity in the presence of both a visual (CI^nothreat^=63-69% vs. CI^threat^=56- 58%, Fig. 2f, Extended Data Fig. 3d-f) or mechanical threat (CI^threat^=42%, Extended Data Fig. 1g). In contrast, optogenetic activation of TRH^R23E12^ in the absence of the threat caused male flies to terminate courtship and exhibit defensive behaviors (CI^lightOFF^=69% vs. CI^lightON^=23%, DI^lightOFF^=31% vs. DI^lightON^=60%, Fig. 2g, Extended Data Fig. 3g). Taken together, these data suggest that TRH^R23E12^ neurons may integrate threats of different modalities in order to act as general effectors of threat responses, irrespective of the specific nature of the threat or the behavior exhibited by the fly.

We next sought to identify the 5-HT receptor(s) involved in inhibiting courtship. *Drosophila* contain five highly conserved 5-HT G-protein-coupled receptors^70^, all of which appear to be expressed in P1 neurons^71^. We individually downregulated each 5-HT receptor in P1 neurons using RNAi^70^. Knocking down 5-HT1A, 5-HT2A or 5-HT1B did not significantly affect threat responses (CI^threat^=8-31%, Extended Data Fig. 3h). However, flies deficient in either 5-HT7 or 5-HT2B in P1 responded less to the threat and showed higher courtship levels than controls (5- HT7: CI^threat^=42%, vs.; 5-HT2B: CI^threat^ =34%, vs.; Controls: CI^threat^=15%, Fig. 2h, Extended Data Fig. 3h). Given that knocking down 5-HT7 gave the strongest phenotype, we focused our analysis on this receptor. In live imaging experiments, we found that applying 5-HT to the brain decreased GCaMP6s fluorescence in P1 neurons (ΔF/F0post=-4.5%, Fig. 2i), an inhibitory effect that was abolished when 5-HT7 expression was decreased in P1 (Fig. 2j, Extended Table 2). These experiments suggest that 5-HT suppresses courtship by inhibiting P1 cells via 5-HT7. However, we do not exclude the possibility that 5-HT2B might also be required in P1 neurons to suppress courtship upon threat detection.

While *Drosophila* 5-HT7 can act as an excitatory receptor by increasing intracellular cAMP^70,72^, the same GPCR can be excitatory or inhibitory depending on the associated G protein and the specific cell type involved^70,73,74^. To investigate the mechanism by which 5- HT7 triggers inhibition, we downregulated different G-proteins and evaluated behavioral responses. Knocking down the inhibitory Gαi protein in P1 neurons prevented males from prioritizing defensive responses over courtship (CI^threat^=51% vs. Control CI^threat^ =26%, Extended Data Fig. 3i) and abolished the inhibitory effect of serotonin on P1 calcium activity (Extended Data Fig. 3j). Knocking down either 5-HT7 or Gαi in P1 neurons did not affect the response of solitary males to the threat (Extended Data Fig. 3k). These findings collectively suggest that, in response to visual threats, 5-HT inhibits P1 via 5-HT7-Gαi signaling, thereby suppressing courtship.

Given that inhibiting P1 suppresses courtship only transiently^17^, we reasoned that other neurons are involved in sustaining the threat-induced inhibition of courtship. One potential candidate is plP10 neurons, which descend from the brain to the wing motor region in the ventral nerve cord and are crucial for eliciting courtship song^63^ (Fig. 2k). Indeed, optogenetic activation of plP10 resulted in high and sustained courtship levels throughout the threat delivery (CI^threat+lightON^=99%, vs. CI^threat+lightON^=7-12% for genetic controls, and CI^threat+lightOFF^=26% for experimental flies, Extended Data Fig. 4a,b), whereas optogenetic inhibition of plP10 via GtACR1 in the absence of the threat robustly suppressed courtship (CI^lightON^=12%, DI^lightON^=12% vs. CI^lightOFF^=50%, DI^lightOFF^=35%, Extended Data Fig. 4c,d).

We next sought to determine whether—as with P1 neurons— plP10-mediated courtship inhibition is serotoninergic. We used RNAi to downregulate each of the *Drosophila* 5-HT receptors in plP10 neurons and assessed the effect on the males’ courtship/escape behavior. Males in which the 5-HT2B receptor was knocked down in plP10 neurons showed similar levels of courtship irrespective of the presence of the threat (CI^nothreat^=40%, CI^threat^=33%, Fig. 2l, Extended Data Fig. 4e). 5-HT2B knockdown in plP10 neurons had no effect on the defensive behaviors *per se* (Extended Data Fig.4f), showing that plP10 is specifically involved in modulating courtship behaviors. Knocking down other 5-HT receptors did not significantly affect the behavioral choice (CI^threat^=4-23%, Extended Data Fig. 4e). In calcium imaging experiments, 5-HT application significantly decreased the activity of plP10 neurons (ΔF/F0 post=-19, Fig. 2m), and this inhibitory effect was abolished by reducing 5-HT2B expression in plP10 (ΔF/F0post=0, Fig. 2n, Extended Data Table 2). Together, these results show that 5-HT inhibits plP10 neurons via 5-HT2B.

Like 5-HT7, 5-HT2B is not considered to be an inhibitory receptor^70^. To elucidate how 5-HT2B can inhibit plP10, we downregulated different G-proteins. Depleting the inhibitory Gαo protein - but not Gɑi - from plP10 neurons prevented males from prioritizing escape responses over courtship (Extended Data Fig. 4g) and abolished the inhibitory effect of serotonin on plP10 calcium activity (Extended Data Fig. 4h). Taken together, these findings suggest that 5-HT2B inhibits plP10 via Gαo.

While direct evidence for connections between 5-HT^PMPD^, P1, and plP10 neurons awaits confirmation, our findings suggest a model whereby, upon threat detection, LC16 neurons activate 5-HT^PMPD^ neurons via an intermediate neuron. The threat-driven release of 5-HT inhibits P1 and plP10 neurons, allowing flies to prioritize survival over sex (Fig. 2o). Notably, threat-driven inhibition of courtship was not fully prevented by blocking serotonergic signaling to either P1 or plP10 individually (Fig. 2h,l), indicating that both neural pathways must be inhibited to halt courtship upon threat detection.

### Courtship progress decreases threat responses

Our findings show that male flies abort courtship in response to threats presented directly after courtship onset. But is this also the case in advanced stages, when they have already invested in courtship and are likely closer to achieving mating? Indeed, previous research has demonstrated that the value of sensory information can be influenced by ongoing behavior and internal state^16,75–77^ . Therefore, to investigate whether male flies respond differently to the visual threat at advanced stages, we delivered the visual threat at progressive courtship stages (‘early’, ‘middle’ and ‘late’ threat, corresponding to 7 s, 2 min and 4 min after courtship initiation, respectively, Fig. 3a, Extended Data Fig.5a,b). We found that as males advanced further in the courtship process, the threat was less effective at making them stop courting (CI^earlythreat^=9%, CI^middlethreat^=19%, CI^Iatethreat^=41%, Fig. 3b) and show defensive behaviors (DI^earlythreat^=58%, DI^middlethreat^=19%, DI^Iatethreat^=0%, Fig. 3c and Extended Data Fig.5a,b). Moreover, copulating male flies completely ignored visual threats, even after sperm transfer (Extended Data Fig. 5c, d) compared to heat shock threats, which do terminate copulation after sperm transfer^9^. These results collectively suggest that as the courtship ritual progresses, males become increasingly unresponsive to visual threats.

**Figure 3:**
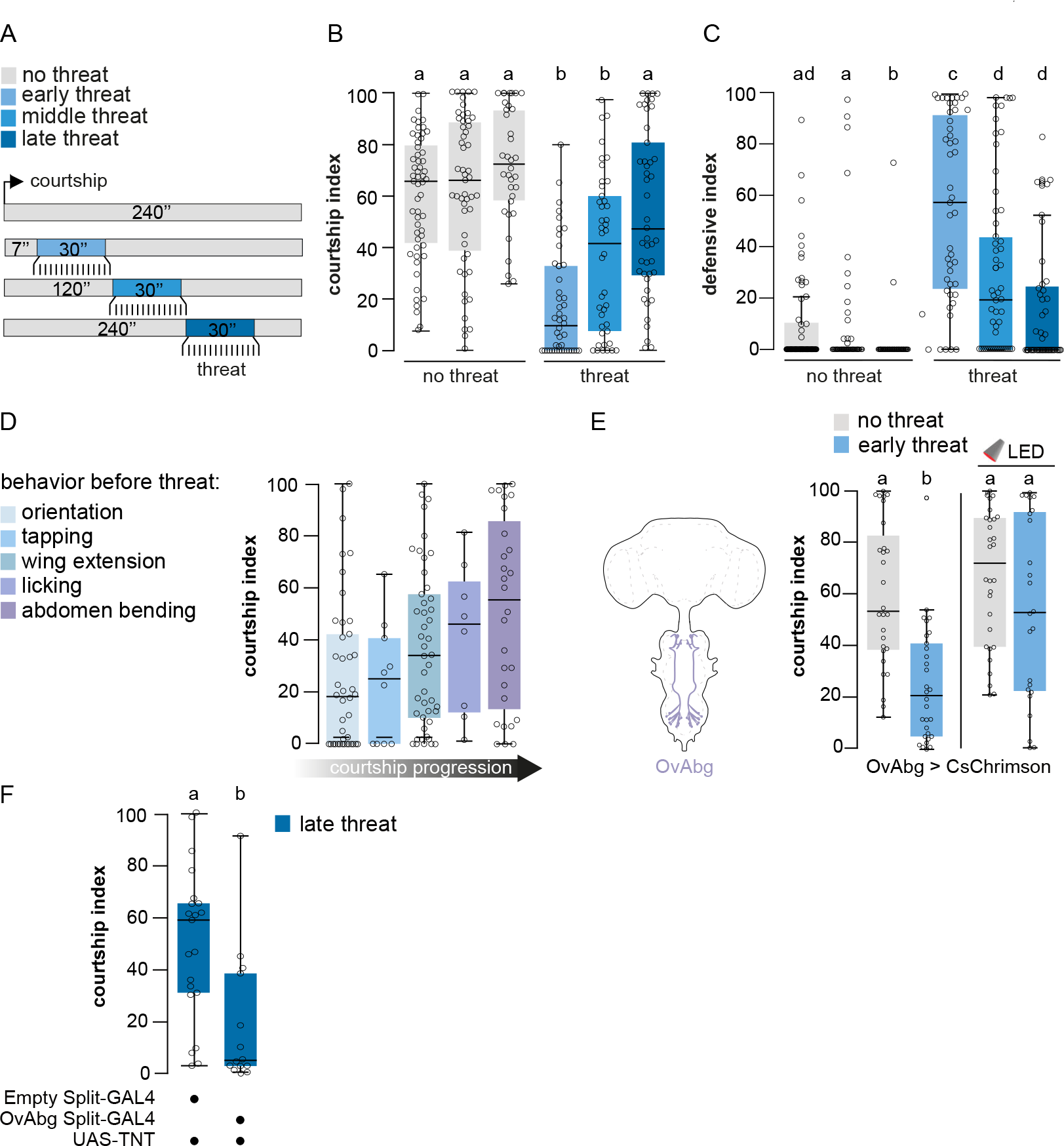
Flies engaged in later courtship steps show reduced threat responses. **(A)** Behavioral protocol: the visual threat is delivered after 7 s (early stage), 120 s (middle stage) or 240 s (late stage) of sustained courtship. Controls for courtship (no threat) have been tested at the same time points in the absence of the threat. **(B)** Courtship index of 5-7 day old male CS in absence of the threat (grey plots, n=48-56) or in presence of an early (30 s), middle (120 s) or late (240 s) threat (blue plots, n=47-52). **(C)** Defensive index of 5-7 day old male CS in absence of the threat (grey plots, n=48-56) or in presence of an early (30 s), middle (120 s) or late (240 s) threat (blue plots, n=47-52). **(D)** Courtship index of CS males from Fig. 3a,b displaying a given behavior before the threat, irrespective of the stage of threat delivery. (orientation n=42, tapping n=10, wing extension n=46, licking n=8 or abdomen bending n=28). **(E)** Left: schematic of OvAbg neurons. Right: courtship index after artificial activation of OvAbg neurons under 660 nm red light using CsChrimson in absence (grey plots, n=15- 19) or presence (blue plots, n=19-20) of the threat. **(F)** Courtship index of males with OvAbg neurons silenced and genetic control in presence of a late threat (240 s, n=14-21).

Female flies signal acceptance and readiness to copulate by ceasing rejection behaviors and slowing down, allowing the male to bend its abdomen and mount them. As a result, abdomen bending may indicate proximity to expected copulation. Thus, we asked whether the observed reduced response to the threat (Fig. 3b-d, Extended Data Fig. 5a,b) was linked to the execution of advanced courtship steps. Indeed, males that actively engaged in abdominal bending before threat presentation were less likely to interrupt courtship, regardless of when the threat was delivered (CI^abd.ben^=59%, Fig. 3d, Supplementary Video 5). In line with this, optogenetically inducing abdomen bending via the activation of a small subset of abdominal ganglion neurons (OvAbg^78^ also diminished threat responses (Fig. 3e). Crucially, silencing OvAbg neurons made flies reduce courtship in response to the threat during late courtship stages, indicating that OvAbg activity is required for late-courtship males to ignore the threat (Fig. 3f).

### Dopamine activity ramps up as courtship progresses

Having established that as males advance in their courtship, they become less responsive to threats, we next explored how courtship progress is integrated within the neural circuitry that arbitrates between courtship and escape. Dopamine is intricately linked with behavior engagement, as well as the perceived value and distance of a reward^31,33,76,79,80^. We therefore speculated that courtship progress in *Drosophila* males might correlate with changes in dopamine neuron activity. To test this notion, we focused on a group of dopamine neurons targeted by TH-C1-GAL4, some of which have been shown to modulate mating drive^22^ (Fig. 4a, Extended Data Fig.7d, Extended Table 3). In behavioral experiments, we found that the activation of TH-C1 neurons during early courtship caused males to ignore threats and continue courting the female (Fig. 4b, Extended Data Fig. 6a,b). Activation of TH-C1 neurons did not affect courtship performance (CI^nothreat+lightON^=60% vs. genetic controls CI^nothreat+lightON^=58- 61%, Fig. 4b) or visual threat detection *per se* (Extended Data Fig. 6c).

**Figure 4:**
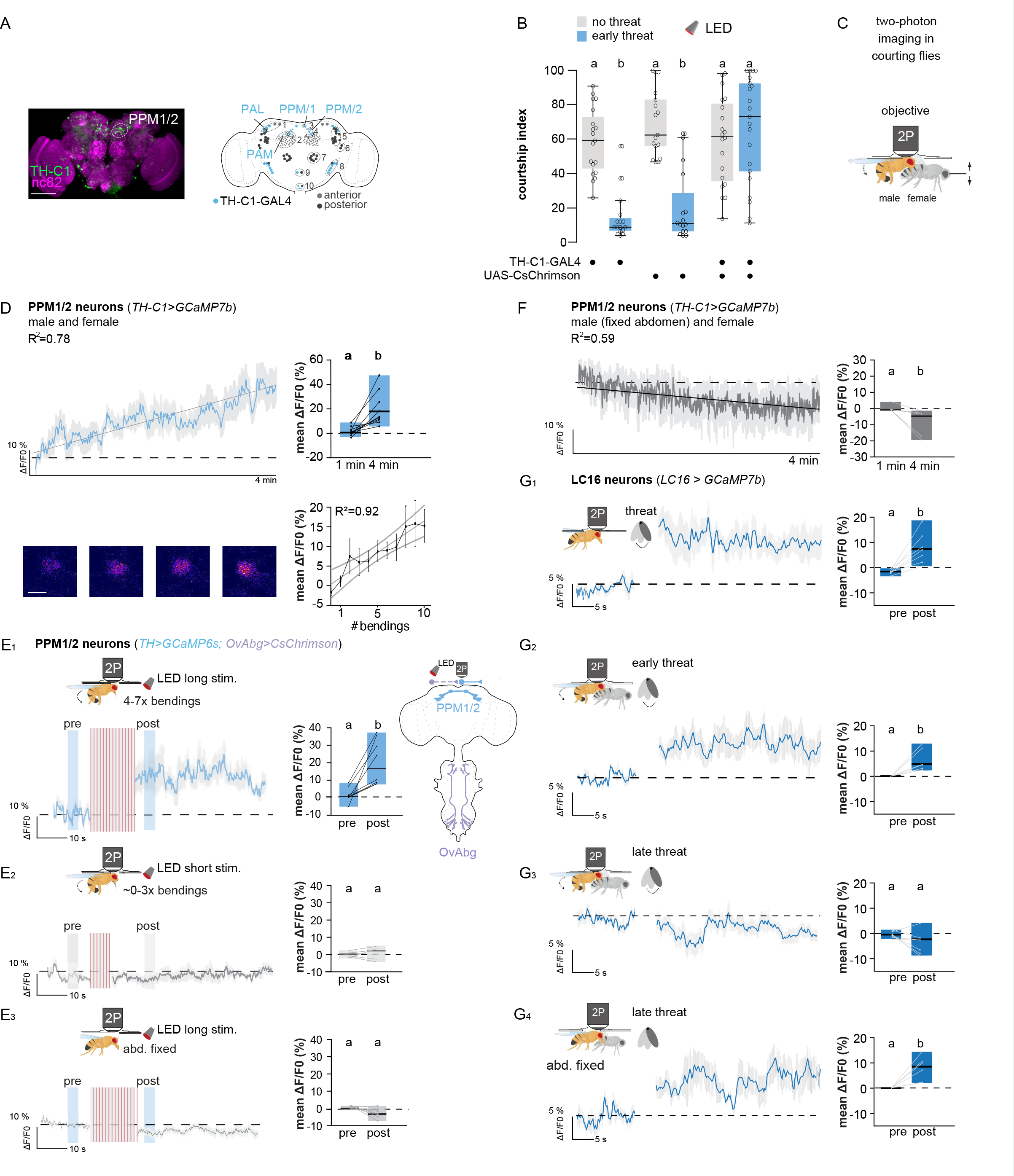
Ramping dopamine release reflects courtship progress. (A) Left panel: UAS-mCD8-GFP (green) expressed under the control of the TH-C1 GAL4 driver in the adult brain; neuropil counterstained with anti-nC82 (purple). Scale bar: 50 μm. Right panel: schematic representation of TH-C1-GAL4 expression pattern (blue) and the different anterior (grey) and posterior (black) dopamine clusters in the adult central brain. 1: PAL (protocerebral anterior lateral), 2: PAM (protocerebral anterior medial), 3: PPM1 (protocerebral posterior medial 1), 4: PPM2 (protocerebral posterior medial 2), 5: PPL1 (posterior protocerebrum lateral 1), 6: PPL2c (posterior protocerebrum lateral 2c), 7: PPM3 (protocerebral posterior medial 3), 8: PPL2ab (posterior protocerebrum lateral 2ab), 9: T1 (thoracic 1), 10: SB (subesophageal zone). (B) Courtship index after artificial activation of TH-C1 neurons under 660 nm red light using CsChrimson in absence (grey bars, n=17-20) or presence (blue bars, n=16-21) of the threat. (C) Schematic of the two-photon live imaging set up in courting flies. The male is tethered to an imaging chamber that allows access of the objective to the brain but leaves the legs, proboscis and abdomen free to move. A decapitated female is tethered to a movable arm that can be approached to the male orienting the female’s abdomen towards the head of the male. Presence of the female results in typical courtship behaviors such as tapping, licking as well as abdomen bending. (D) Upper left: fluorescence intensity trace (ΔF/F0) over 4 min in male flies expressing GCaMP7b in PPM1/2 neurons while exposed to a female fly under the microscope. Upper right: quantification of mean ΔF/F0 during 1^st^ min compared to 4^th^ min (20 s time windows each, starting at 0 s and 220 s) (n=10). Bottom left: shows representative fluorescence heatmap of PPM1/2 neurons at timepoint 1 min, 2 min, 3 min and 4 min. Scale bar: 2µm. Bottom right: correlation of calcium activity and number of abdominal bendings. **(E1-3)** Fluorescence intensity trace (ΔF/F0) over time pre and post long or short optogenetic stimulation of OvAbg neurons under 590 nm red light in male flies expressing GCaMP6f in PPM1/2 neurons with the abdomen either free or fixed. Right: Quantification of mean ΔF/F0 comparing pre and post stimulation as indicated by blue squares (n=8). Please see Extended Data Figure 6m for further details on the LED protocol for optogenetic activation of OvAbg. **(F)** Fluorescence intensity trace (ΔF/F0) over 4 min in male flies expressing GCaMP7b in PPM1/2 with a fixed unmovable abdomen while paired with a female fly under the microscope. Right: Quantification of mean ΔF/F0 during 1^st^ min compared to 4^th^ min (20 s time windows each) (n=6). **(G1-4)** Fluorescence intensity trace (ΔF/F0) over time of male flies expressing GCaMP7b in LC16 neurons pre and post 30 s of threat exposure under the microscope (1) male alone (2) together with a female in early threat condition presented after 10 s (3) together with a female in late threat condition presented after 4 min and (4) together with a female in late threat condition presented after 4 min with a fixed unmovable abdomen. Right: Quantification of mean ΔF/F0 comparing the 30 s pre and post time windows (n=6-7).

We thus sought to test whether TH-C1 dopamine neural activity might change with courtship progress. To this end, we developed a calcium imaging system for tracking neural activity as a male fly progresses through courtship under a two-photon microscope (Fig. 4c, Extended Data Fig. 6d,e, Supplementary Video 6). When exposed to a virgin immobile female, tethered males displayed early and late courtship steps (e.g., tapping, licking and abdomen bending). During the final tier (161-240 s) of the 4 min experimental time window, males showed an increase in late courtship actions, such as abdomen bending (Extended Data Fig. 6d,e, Supplementary Video 6). Remarkably, GCaMP7b recordings revealed a gradual increase in calcium signal within a group of ∼7 dopamine neurons per hemisphere, named PPM1/2 (protocerebral posterior medial 1/2) during courtship progress (Fig. 4d). The mean normalized GCaMP7b fluorescence increased by ∼17% after 4 min when compared to the signal in the first minute (R^2^=0.78, Fig. 4d). This effect was absent when the focal male was paired with another male, which elicited minimal abdomen bending events (R^2^=0.29, Extended Data Fig. 6f, Extended Table 2). Importantly, the dopamine signal returned to baseline when the female was moved away (Extended Data Fig. 6g). In contrast to ramping PPM1/2 neurons, GCaMP7b signal did not increase in adjacent cell bodies located within the same focal plane (Extended Data Fig. 6g). Similarly, neither TH-C1+ PAL dopamine neurons (linked to mating drive^22^ nor TH-C1+ PAM dopamine neurons (involved in courtship reward^81^ increased their activity as males progressed through courtship (Extended Data Fig. 6h,i, Extended Table 2). Together, these findings suggest that the calcium ramping observed in PPM1/2 neurons is specific to courtship progression.

In line with our behavioral findings, we observed a correlation between the display of abdomen bending and an increase in the GCaMP7b signal in PPM1/2 neurons during live calcium imaging (Fig. 4d bottom right panel, Extended Data Fig. 6j), suggesting a direct link between abdomen bending and dopamine neural activity. To confirm that abdomen bending - and not other courtship actions - drives PPM1/2 ramping, we measured calcium signals in PPM1/2 neurons over 4 minutes in males with their proboscis or legs immobilized, rendering them unable to lick or tap the female (Extended Data Fig. 6k,l; Supplementary Video 7-8, Extended Table 2; note that males do not display wing vibrations in our setup, see *methods* for details). Stimulation of abdomen bending via OvAbg in solitary males led to a ∼20% increase in PPM1/2 activity (Fig. 4e1). This effect was dose-dependent, as dopamine ramping was observed after prolonged LED stimulation (Fig. 4e1,2, and Extended Data Fig. 6m for details on protocol), which elicited 4-7 bending events (Extended Data Fig. 6m) but not short LED stimulations, which elicited 0-3 bending events (Extended Data Fig. 6m, Extended Table 2). Notably, males exhibited abdominal bending events even after the long LED stimulation ceased, suggesting that this ongoing behavior may sustain elevated PPM1/2 calcium levels (Extended Data Fig. 6m). Hence, our findings support the conclusion that PPM1/2 dopamine ramping is primarily driven by abdomen bending during courtship progression.

The dopamine activity ramping could be driven by either sensory information triggered by the abdomen bending action (proprioception) or the predictive signal in anticipation of this movement (efferent copy). To address this, we immobilized the male’s abdomen using glue, which prevented males from bending it while preserving other courtship behaviors such as tapping and licking (Fig. 4f, Extended Data Fig. 6n). Following this manipulation, the GCaMP7b signal in PPM1/2 neurons did not show the expected ramping behavior. Instead, it decreased significantly over time (ΔF/F0=-7%, R2=0.53; Fig. 4f, Extended Table 2), suggesting that proprioceptive feedback from abdomen bending, rather than efferent copy from a command circuit, is required to ramp up dopamine activity in PPM1/2 neurons. Consistent with these findings, dopamine ramping triggered by optogenetic activation of OvAbg neurons only occurred when solitary males were physically able to bend their abdomen (Fig. 4e3, Extended Table 2). These findings indicate that abdomen bending during late stages of courtship suppresses LC16 visual threat responses. Supporting this observation, in live imaging experiments, we found that while LC16 neurons responded to threats during early courtship (Fig. 4g1-2), these responses were absent in the late courtship stages (Fig. 4g3). Notably, when abdomen bending was mechanically inhibited, LC16 threat responses during late courtship were restored (Fig. 4g4). Together, our findings support a role for abdomen bending in modulating dopaminergic activity levels, which integrate proprioceptive feedback and ultimately induce a late courtship state.

### Copulation attempts drive dopaminergic inhibition of the visual threat pathway

Our findings indicate that dopamine activity ramps up with copulation attempts (Fig. 4d), and engagement in abdomen bending correlates with impaired threat responses (Fig. 3d-e). We therefore investigated how increased dopamine levels might translate courtship progress into reduced threat responses. Utilizing the female connectome, we found that LC16 neurons receive direct input from PPM1/2 at their axon terminals but not from other dopaminergic clusters (Fig. 5a). Thus, we hypothesized that PPM1/2 neurons directly modulate the perception of visual threats.

**Figure 5:**
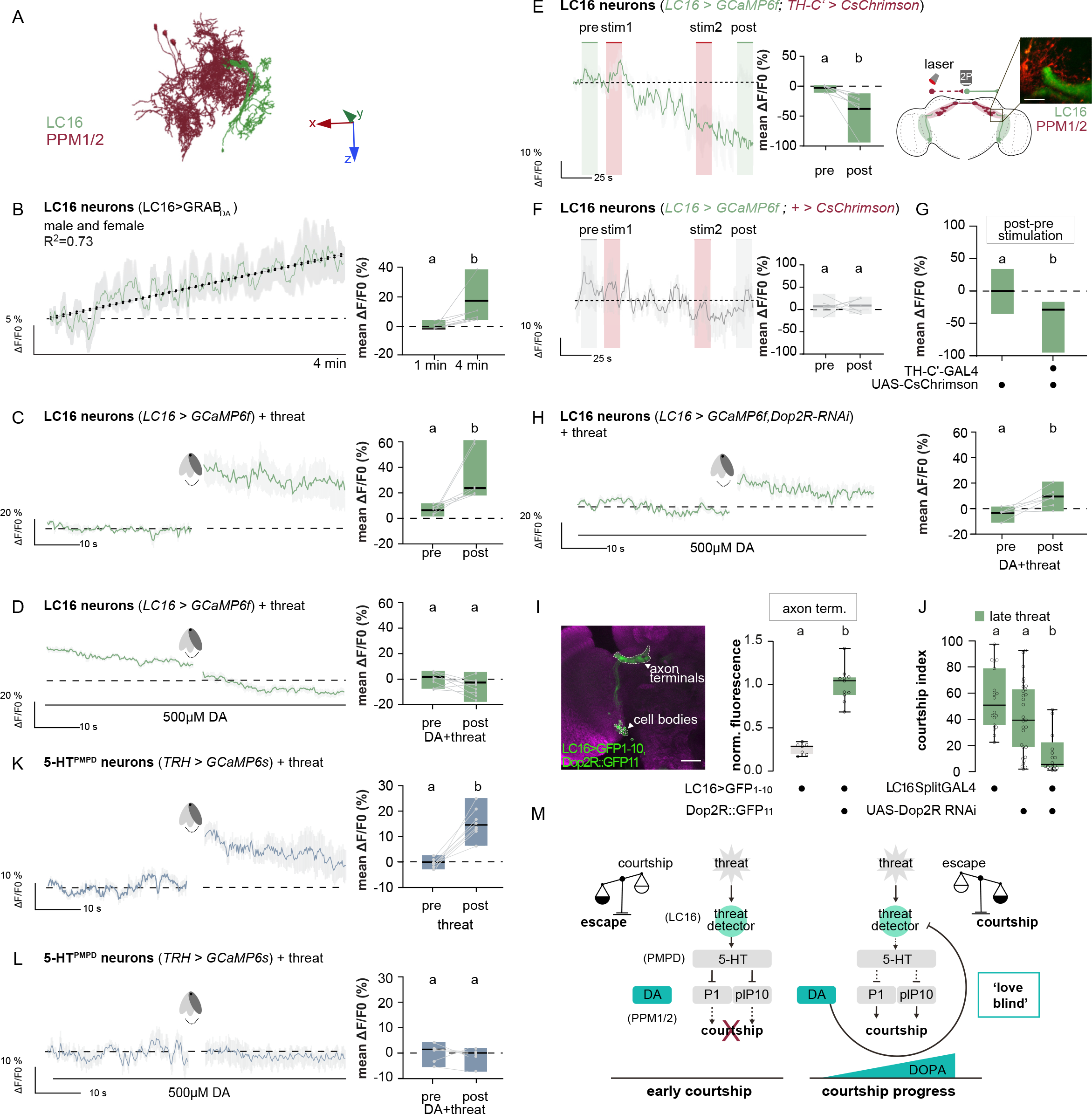
Ramping dopamine inhibits the visual threat pathway. **(A)** Electron microscope reconstruction images of LC16 neurons (green) and PPM1/2 (PPM1201-03) dopaminergic neurons (red) in the female central nervous system. x,y,z arrows represent the anterior/posterior, medial/lateral and dorsal/ventral axis, respectively. **(B)** Left: Fluorescence intensity trace (ΔF/F0) over 4 min in male flies expressing the GRABDA sensor in LC16 neurons when paired with a female fly under the microscope. Right: Quantification of mean fluorescence intensity during 1^st^ min compared to 4^th^ min (20 s time windows each) (n=7). **(C-D)** Fluorescence intensity trace (ΔF/F0) over time of male flies expressing GCaMP6f in LC16 neurons pre and post **(C)** 30 s of threat exposure under the microscope; **(D)** threat exposure in presence of 500 µM dopamine. Right: Quantification of mean ΔF/F0 comparing the 30 s pre and post time windows (n=7-9). (E) Left: Fluorescence intensity trace (ΔF/F0) over time including 2 x 15 s of PPM1/2 neuron activation using CsChrimson under 1040 nm two-photon laser stimulation in male flies expressing GCaMP6f in LC16 neurons. Middle: Quantification of mean ΔF/F0 comparing pre and post stimulation windows as indicated by green squares (n=8). Right: Schematic illustrating GCaMP expression in LC16 neurons and CsChrimson expression in PPM1/2 neurons, and image showing imaging region (Scale bar: 15µm). (F) Fluorescence intensity trace (ΔF/F0) over time including 2 x 15 s of 1040 nm two-photon laser stimulation in male flies expressing GCaMP6f in LC16 neurons without CsChrimson. Right: Quantification of mean ΔF/F0 comparing pre and post stimulation windows as indicated by grey rectangles (n=6). (G) Comparison between mean ΔF/F0 (post-pre) of flies expressing GCaMP6f in LC16 neurons with and without CsChrimson in PPM1/2 neurons. (H) See C-D with additional Dop2R knock down in LC16 neurons. Right: Quantification of mean ΔF/F0 comparing the 30 s pre and post time windows (n=7-9). (I) Left: UAS-spGFP1-10 expressed under the control of LC16 split-GAL4 combined with tagging endogenous Dop2R with spGFP11 (Dop2R::GFP11). Neuropil counterstained with anti-brp (nc82; magenta). Dashed white outlines represent the regions of interest for the axon terminals and cell bodies of the LC16 neurons. Right: quantified GFP fluorescence in LC16 axon terminals in flies with UAS-spGFP1-10 expressed under the control of LC16 Split-GAL4, with or without Dop2R::GFP11. GFP fluorescence was normalised to the average fluorescence in the LC16 axon terminals in LC16>GFP1-10, Dop2R::GFP11 flies (n=7-11 brains, scale bar: 25 μm). (J) Courtship index of males with the Dop2R receptor knocked down specifically in LC16 visual projection neurons using RNA interference, and genetic controls in presence of a late threat (240 s, n=18-29). (K) Fluorescence intensity trace (ΔF/F0) over time of male flies expressing GCaMP6f in 5- HT^PMPD^ neurons pre and post 30 s of threat exposure under the microscope. Right: Quantification of mean ΔF/F0 comparing the 30 s pre and post time windows (n=8). (L) Fluorescence intensity trace (ΔF/F0) over time of male flies expressing GCaMP6f in 5- HT^PMPD^ neurons pre and post 30 s of threat exposure in presence of 500 µM dopamine. Right: Quantification of mean ΔF/F0 comparing the 30 s pre and post time windows (n=6). (M) Trade-off between reproduction and survival working model. Solid and dashed lines indicate direct and indirect connections respectively.

To functionally test the PPM1/2-to-LC16 connection, we tracked dopamine release onto LC16 presynaptic terminals in live imaging experiments, using a fluorescent G-protein-coupled dopamine receptor-activation-based (GRABDA) sensor^82^. We found a steady increase in GRABDA fluorescence in males paired with a female, indicating a gradual increase in dopamine release onto LC16 presynaptic terminals (ΔF/F0 4 min=+8%, R^2^=0.73, Fig. 5b), consistent with the steady increase in PPM1/2 activity (Fig. 4d). This dopamine ramping profile was not observed in males paired with another male (ΔF/F0 4 min=-3%, R^2^=0.32, Extended Data Fig. 7a, Extended Table 2). These findings suggest that the gradual release of dopamine onto LC16 may contribute to the reduced responsiveness of LC16 axonal terminals to visual threats during courtship progress.

To directly test whether dopamine shuts down visual threat responses in LC16 neurons, we recorded activity in LC16 presynaptic terminals during threat delivery while simultaneously administering dopamine. Not only did application of dopamine reduce LC16 baseline calcium activity (ΔF/F0=-5%, Extended Data Fig. 7b) but, more importantly, it completely suppressed the threat-driven LC16 response (ΔF/F0=+20% threat alone, ΔF/F0=0, threat + dopamine, Fig. 5c,d, Extended Table 2). Moreover, intra-vital focal injection of dopamine using a micropipette directly onto LC16 presynaptic terminals caused a robust decrease in LC16 calcium activity, suggesting axo-axonal modulation of dopamine at the LC16 output level (ΔF/F^dopamine^= -15% vs. ΔF/F^saline^=-2%, Extended Data Fig. 7c, Extended Table 2). Also, optogenetically stimulating PPM1/2 dopamine neurons did not cause a time locked effect but rather a gradual decrease in GCaMP6s fluorescence in LC16 neurons (Fig. 5e-g, Extended Table 2) and – consistent with dopamine administration – reduced threat responses in LC16 (ΔF/F0=+4% threat alone, Extended Figure 7e1;ΔF/F0=-2,threat + optogenetic PPM1/2 activation, Extended Figure 7e2; ΔF/F0=+10% threat + LED without CsChrimson, Extended Figure 7e3, Extended Table 2). This dopamine-induced inhibition seems to act through dopamine D2-like receptors (Dop2R), since expressing Dop2R-RNAi in LC16 neurons partially rescued the threat response and prevented LC16 inhibition by focal dopamine injection (ΔF/F0=+10%, Fig. 5h; ΔF/F^dopamine^= -15% vs. ΔF/F^dopamine+DOP2R^ RNAi=-2%, Extended Data Fig. 7c). Dop2R expression in LC16 was confirmed using reconstitution of split-GFP where we tagged endogenous Dop2R with spGFP11 and expressed cytoplasmic spGFP1-10 under LC16 split-GAL4^83^ (Fig. 5i, Extended Fig. 7f). Reconstituted GFP signal was significantly higher than in split-GFP1-10-only controls in LC16 presynapses, but not LC16 cell bodies (Fig. 5i, Extended Fig. 7f), indicating that Dop2R receptors are localized in the axon terminals, proximal to PPM1/2 neurons. Crucially, when flies expressing Dop2R-RNAi in LC16 neurons were tested in the behavioral assay, their behavior shifted from courtship to defensive responses during late courtship stages (CI^latethreat^=4.5% vs. CI^latethreat^=39-51% for genetic controls, Fig. 5j).

Our findings suggest that dopaminergic modulation of outgoing activity from LC16 neurons modulates 5-HT^PMPD^ neurons to the visual threat. Supporting this, we found that while 5- HT^PMPD^ activity increased following threat delivery (Fig. 5k), this response was abolished upon the addition of dopamine (Fig. 5l, Extended Table 2). Moreover, this dopamine-driven inhibition of 5-HT^PMPD^ threat responses was in turn blocked by knocking down Dop2R in LC16-GAL4 neurons (Extended Data Fig. 7g).

Together, our findings suggest that dopamine signalling from PPM1/2 acts through Dop2R to shut down LC16-mediated threat detection, allowing males to persist in courtship despite the presence of a threat.

## DISCUSSION

Amorous adventures can lead us to pursue risky actions. This ’love blindness’ has not only frequently been described in literature^84^ but is indeed the product of a fundamental function of life: weighing up risks against opportunities. Our study showcases this balancing act by demonstrating that imminent mating success disrupts threat perception in *Drosophila* males, rendering them ‘love-blind’. We show that, at the first sign of a visual threat, LC16 visual neurons trigger serotonin-mediated inhibition of key courtship nodes (P1 and plP10), prompting flies to abort courtship (Fig. 2o). But as courtship progresses (as reported by abdomen bending), the activity within dopamine neurons (PPM1/2) gradually increases (Fig. 4d). PPM1/2 activity in turn suppresses LC16 activity, allowing flies to persist in courtship and ignore external threats when close to mating (Fig. 5m). Thus, risk-benefit arbitration is dynamically modulated by goal proximity and is under dopaminergic control.

What is more evolutionarily advantageous: escaping threats, or attempting to copulate in dangerous situations? The trade-off lies in the fact that individuals must take risks to reproduce, while also needing to survive to achieve mating^85^. Examples of risk/reward trade-offs abound in the animal kingdom. Animals become less risk-averse when the opportunity cost of dying (foregone future mating opportunities) is lower. For example, late in the breeding season when future reproductive chances are limited, stickleback fish take the risk of displaying bright coloration despite the presence of predators^86^. Furthermore, animals become less risk-averse when the expected reproductive rewards are higher. Male mice, for instance, become less afraid of predator odors after smelling estrous female odors^87^, while male moths following pheromone plumes ignore ultrasound cues that simulate an approaching bat^88^. Our results now provide a mechanistic framework for how such processes can occur in the brain and how risk vs. reward trade-offs are calculated based on mating success. Future studies should consider how courtship progress is integrated with external factors, physiological state, and mating history to gain a deeper understanding of the trade-off between sex and survival and its broader implications for life-history evolution.

Our study suggests that abdomen bending triggers a late courtship state via proprioceptive feedback ramping up dopaminergic activity. While the proprioceptive neurons involved remain unknown, the role of interoceptive and exteroceptive sensory feedback is well-established in *Drosophila* reproductive behaviors, including male copulation^89^, female receptivity^90^ and egg- laying^91^. Interestingly, PPM1/2 neurons do not respond to discrete abdominal bending events with phasic, time-locked responses. Instead, the observed gradual ramping of PPM1/2 activity suggests that PPM1/2 neurons integrate proprioceptive inputs over time, leading to a gradual increase in tonic calcium levels. We propose that this gradual rise in calcium activity within dopaminergic neurons stems from the continued integration of both female sensory cues and proprioceptive feedback from abdominal bending. It will be interesting in future studies to test candidate biophysical mechanisms underlying this integration, such as neuromodulatory regulation of spontaneous activity and intrinsic excitability^92–94^.

In addition to dopamine ramping suppressing threat detection, other parallel modulatory mechanisms might work together to prioritize courtship when copulation is imminent. Indeed, PPM1/2 activation prevents threat responses in courting males but not solitary males, suggesting that PPM1/2 inhibits but does not entirely silence LC16 output, such that the reduced output can still drive escape behavior in solitary males but cannot outcompete courtship drive. Other parallel mechanisms might reduce serotonergic threat responses or reduce the sensitivity of central courtship nodes to serotonergic inhibition in conourting males. For example, sexual arousal due to increased P1 activity gates the perception of female-related visual cues during courtship^16^, and after courtship begins, P1 activity is thought to be sustained by recurrent activation, facilitated by dopamine released from neurons in the anterior superior medial protocerebrum (SMPa) ^17,22,64^ (though P1 activity does not ramp up the way PPM1/2 neurons do) ^57^. Future experiments should test if this recurrent activity is facilitated by the same PPM1/2 neurons that ramp up as courtship progresses.

While this study focused on male *Drosophila* threat responses during courtship, an interesting question pertains to whether females react similarly. Unlike their male counterparts, females do not show a distinct progressive behavioral pattern during courtship, but they do signal receptivity to copulation by slowing down and opening their vaginal plates^20^. Studies have described a role for dopamine in modulating female sexual receptivity^81,95^ ; however, the link between dopamine and acceptance behavior during courtship, and its subsequent impact on threat responses, remains an open question. As females need to survive to produce offspring, they might have greater incentive to escape threats than courting males. One possibility is that PPM1/2 dopamine ramping is not present in courted females, leading them to consistently prioritize survival over reproduction.

Studies in mammals have reported dopamine ramping in diverse behavioral trials leading to reward, including goal-directed navigation and multi-step tasks^33,79,80,96,97^. Such ramping release profiles have been proposed to supply the motivational drive required to sustain goal pursuit, as they scale with distance to reward^33,79,80,96–98^. Our findings challenge the notion that build-up of dopamine activity primarily encodes reward expectation^99–101^. Instead, our study reveals a novel function of dopamine ramping as a gradual sensory filter system, which shuts down the visual threat pathway as courtship progresses. Such a mechanism would render animals in a ‘love blind’ state, allowing them to pursue their reproductive goal despite the danger.

Dopaminergic modulation of sensory signaling has been shown in different species. For instance, in lampreys and zebrafish, dopamine neurons modulate responses to visual features like looming cues^50,102^. In rodents, dopamine has been demonstrated to influence subcortical responses to unexpected auditory stimuli^103^. In humans, antipsychotic drugs are thought to act on D2 receptors^104^, the mammalian homolog of Dop2R, suggesting that D2-like receptors may have a common role in top-down regulation of sensory perception, whether in generating hallucinations or ignoring visual threats. Given the striking similarities in the cellular biology of dopamine neurons across species^76,103,105–107^, dopamine-mediated filter systems may be a general mechanism for blocking sensory cues that compete with more important goals.

## Methods

### Material and Methods

#### Resource availability

Requests for additional information and reagents should be addressed to the lead contact Carolina Rezaval. All data generated in this paper can be shared upon request.

#### Fly husbandry and strains

Flies were reared at 25°C or 30°C for RNAi experiments, with 40-50% humidity on a standard cornmeal-agar food detailed below in a 12-hr light:dark cycle. Canton-S (CS) strain flies were used as wild-type. Additional strains used and their sources are outlined in Table 3. Flies were sorted under CO_2_ anaesthesia within 6 hours of emergence and housed in same-sex groups of 20, except for the males to be tested in behavior, which were kept in groups of 4 per vial. Virgin females for behavior were collected using the *hs-hid* conditional virginator transgenic line. L3 larvae were heat shocked at 37°C for 1.5 hours.

| Ingredient | Quantity | Supplier and catalog number |
| --- | --- | --- |
| water | 1300ml |  |
| agar | 17g | Fisher Scientific 11954041 |
| 10% nipagin diluted in ethanol | 50ml | Stratech Scientific Ltd - T2300 |
| dextrose | 150g | Merck Life Science Limited - PHR1000-1G |
| yeast | 40g | Scientific Laboratory Supplies - FLY1132 |
| cornmeal | 170g | Scientific Laboratory Supplies FLY1110 |
| water (for mixing) | 400ml |  |

#### Trans retinal food

All *trans* retinal (R2500-100MG Sigma-Aldrich, #CAS number: 116-31-4) was stored at –20 °C as a 50 mM stock solution diluted in ethanol and wrapped in foil. To blend retinal homogeneously into the food, 60 μl of stock solution was directly pipetted into 6 ml vials of liquid cornmeal-yeast food except for the experiment Fig. 3e where OvAbg flies where not exposed to food supplemented with trans retinal factor.

### Behavior

#### Threat set-up

Experiments were recorded at 27 frames per second using a Mako U-130B camera mounted with and IR filter (MIDOPT #BP735-40.5). The visual threat was generated by repeatedly passing a 13cm x 6cm x 2cm 3D printed oblong and opaque paddle through a blue light beam (455 nm). This created an overhead shadow at periodic intervals of 0.3 Hz for 30 s. The paddle was set 5 cm above the courtship chambers (⌀ 20mm) at a 90° angle on a servo motor controlled by an Arduino custom-built code to control the movement parameters of the paddle (frequency (0.3 Hz) and number of cycles (9)). The mechanical threat was generated using a Sony XP500-X speaker playing a loud 3 Hz binaural beat (https://www.youtube.com/watch?v=Y-urmCRs61I&t=713s) causing surface vibrations. Courtship chambers were illuminated from the bottom using an infrared backlight.

#### Behavioral assays

Behavioral assays were conducted at 25°C under continuous blue light between 9am and 1pm. Tested males were aged 5-7 day old and transferred to fresh food vials one day before experiments. For males used in optogenetic assays, flies were transferred to food enriched with *all*-trans retinal three days prior to the experiment. Vials containing the retinal factor were wrapped in foil. Virgin females were decapitated and used within a maximum of three hours to preserve chemical signature and motor reflexes during the experiment.

#### Action selection assay

The action selection assay presented a naive male coupled with a decapitated *hs-hid* virgin female with a choice between continuing to court the female or interrupting the ritual in response to the threat. The threat was delivered after consistent courtship of at least 7 s (early), 2 min (middle) or 4 min (late). Only males that started to court during the first 5 min of the trial and until threat delivery were considered in the analysis. All assays were manually analysed using the behavioral analysis software BORIS^109^ and the following parameters were quantified in order to measure the effect of the threat on male courtship behavior:

**Courtship index** – is defined by the percentage of time the male spends courting the female (s) over the total time of the threat delivery (30 s). We considered that males initiated courtship by demonstrating full wing extension and a persistent courtship behavior of at least 7 s toward the female. We considered as courtship the display of stereotyped courtship events that include tapping of the female with the male’s forelegs, licking by showing a proboscis extension toward the female, singing (wing extension and vibration), and attempts at copulation were the male bends the abdomen towards the female and attempts to or mount her. See Fig. 1a for schematical representation of these behaviours.

**Defensive index** – is defined by the percentage of time the male spends displaying defensive behaviors (i.e., escaping and freezing) ^47,110^ (s) over the total time of the threat delivery (30 s).

As a control, the behavior was assessed in the absence of the visual threat during the same time window according to the same criteria.

#### Optogenetic assay

Flies were tested in a transparent circular chamber (⌀20mm, h=5mm for courtship, ⌀24mm, h=3mm for locomotion assay) and illuminated from underneath with either 660 nm (red) or 515 nm (green) light in absence or presence of the threat. Refer to Table 4 for the optogenetic experimental conditions corresponding to each figure. The light was turned ON 1 s before the first threat passed.

#### Locomotion assay

Individual flies were introduced into a circular chamber (⌀24mm, h=3mm) and left to acclimatise for 3 min. After the acclimatation period, flies were subjected to the threat (9 cycles, frequency: 0.3 Hz). The walking speed of the flies (thresholded at values larger than 4 mm/s to be considered as ‘walking’) was assessed using the Ethovision software (https://www.noldus.com/ethovision-xt). The change in walking speed was calculated by subtracting the average walking speed 30 s after from before the delivery of the threat.

#### Two-photon functional imaging

Tethered male flies (3–6 days old) had their head capsules dissected in a sugar-free HL3-like saline-filled imaging chamber with a central hole (for details on fly dissection see ^111^). Flies were then placed under a multiphoton microscope (Femto2D-Resonant by Femtonics Ltd., Hungary), and expressed either the calcium indicator GCaMP or GRABDA in different sets of neurons (see Extended Data Table 3 for details on genotypes). Fluorescence was centred on 920 nm generated by a Ti-Sapphire laser (Chameleon Ultra II, Coherent, CA, USA). Images with a pixel size of 0.3 × 0.3 μm were acquired with a 20x, 1.0 NA water-immersion objective, controlled by MESc v3.5 software (Femtonics Ltd., Hungary). Fast recordings were taken at a speed of 30 Hz with a resonant scan head using mesc software (Femtonics). Analysis was performed using NOSA software (neuro-optical signal analysis) ^112^ and a customized R script. ROIs were manually drawn for analysis. Data was converted into tiff files and processed using a Savitzky-Golay filter or moving average of 2 s when brain movement was strong (e.g. Fig. 4d, 5b). No baseline/photobleaching correction was applied to any of the imaging data. The final time resolution was 6 fps (all Femtonics microscope data) or 2fps (Optogenetic data from Nikon microscope). Mean intensity values were calculated as ΔF/F0 (in %), while F0 was defined as the mean F from baseline activity (first 30 s in Fig. 2i, j, m, n; 4e1-3; 5c, d, h, k, l; Ext. data Fig 3j; 4h; 7b, e1-3. First 20 s in Fig. 4d, f; 5b; Ext. data Fig. 6f, g, h, i, k, l; 7a. First 15 s in Fig.5e, f; Ext. Data Fig. 7e and g. First 2 s in Fig. 2e; Ext. data Fig. 3b.)

#### Threat delivery under the two-photon microscope

The threat was delivered as previously described (see Threat set-up section). The paddle and light source were placed below the microscope and inclined toward the chamber in a way that the passing shadow reached the tethered fly’s eye. Calcium signals in LC16 axons were recorded for 30 s before and immediately after the threat exposure. Since LC16 neurons respond to laser onset, the first 2 s of each recording were excluded from the analysis. Conditions under the microscope were set to >20°C and 40% humidity.

#### Application of serotonin or dopamine

100 µl of serotonin (Sigma Aldrich Cat# H9523) or dopamine (Sigma Aldrich Cat# H8502) diluted in sugar-free HL3 solution was applied directly onto the *Drosophila* brain through the open head capsule. The final concentration was 100 µM for serotonin and 500 µM for dopamine. Calcium signals were recorded 50 s before and immediately after application for 100 s (first 30 s of pre and last 30 s of post were taken for quantification).

#### Courtship progression under the microscope

For examining courtship progression, 5-8 day old virgin male flies were used. Flies were tethered and dissected as previously described, leaving legs and proboscis freely movable (or fixed dependent on the experiment indicated for each figure). Note that the fixation position of the male onto the imaging chamber does not allow for wing extension. Agitated males that did not stop moving for 10 s during the first 5 min under the microscope were discarded. Immediately upon recording initiation, a decapitated 3-5 day old virgin female tethered onto a movable arm controlled by a micromanipulator was presented to the male with her abdomen oriented towards the head of the male fly. Following male contact with the female, calcium or GRABDA signals were recorded for a total duration of 4 min, while the fly behavior was simultaneously observed using a video camera (Thorlabs C1285R12M and SM1D12D iris diaphragm) recording at 7 fps. Abdomen bending was manually analyzed frame by frame. Since tethered flies show typical behavior that includes moving the abdomen back and forth, only full bending events (abdomen bending underneath the thorax) that lasted longer than 5 frames were considered as part of courtship behavior.

#### Optogenetic experiments during in vivo calcium imaging

Experiments were conducted using a Nikon A1R+ multiphoton microscope with a galvo scanner at a speed of 2 Hz. We used the microscope’s two-photon 1040 nm red laser to activate CsChrimson while simultaneously recording the calcium activity within the region of interest (see details for conditions in legends and Extended Data Table 4). To activate OvAbg neurons, experiments were carried out using a Femtonics microscope with the same imaging parameters mentioned previously. A 590 nm LED positioned below and towards the tethered fly was employed for optogenetic activation of CsChrimson (15 or 7 repetitions of 1 s LED-on and 1 s intervals), while recording simultaneously. To activate PPM1/2 neurons during threat delivery, 15 repetitions of red light were used overlapping the 30 s of threat exposure under the microscope. LED stimulation artifacts were removed for clarity. Since the acquisition was carried out continuously, the post sequence shown in the graph displays the fluorescence intensity immediately after the LED stopped.

#### Focal dopamine injection

Fly preparation and imaging was conducted as described before using a Nikon A1R+ multiphoton microscope. The sugar-free HL3-like saline was added with 30 units of Papain (Roche) and applied to the head capsule for 10 min to digest the brain’s glial sheath and facilitate removal. Flies were subjected to local dopamine (10 mM diluted in saline) or saline injection via a micropipette (saline used for injection contained no CaCl2 or MgCl2). The injection solution was labelled with Texas Red (Invitrogen by Thermo Fisher Scientific, dextran, 10,000 MW) to visualize the pipette and the localization of the injections. Multiple (2- 5) injections were given per experiment and averaged, resulting in a single average trace per experiment. Fluorescence traces were extracted using FIJI (ImageJ). F0 for the ΔF/F calculations was the average baseline fluorescence of the 10 frames immediately preceding the injection. Mean area under the curve was calculated using GraphPad Prism. Area under the curve values are presented as mean AUC ± SEM. ROIs were selected manually.

#### Immunohistochemistry

3-5-day old male fly brains were dissected in ice-cold PBS and fixed in 4% paraformaldehyde solution at room temperature for 20 min. Fixed brains were then washed four times in PBST (0.5%) for 30 min and blocked with normal goat serum (5%) for 30-60 min. The brains were then incubated with primary antibodies (anti-GFP chicken Abcam 1:1000 or 1:2000, Cat#13970; anti-dsRed rabbit Takara 1:250, cat#632496; nc82 anti-Brp, DSHB, 1:50) for 2-3 days at 4℃. After 4 x 20 min washes in PBST, the brains were incubated with secondary antibodies (Alexa Fluor 488 goat anti-chicken IgG ThermoFisher Scientific 1:1000 (Cat#A28175 or A32931 1:2000), Alexa Fluor 546 goat anti-mouse, 1:2000, ThermoFisher A11018, Alexa Fluor 546 goat anti-rabbit, 1:2000 ThermoFisher A11071) overnight. After 4 x 20 min washes in PBST, brains were mounted in Vectashield on a glass slide before scanning with a Leica SP8 confocal microscope, a Nikon A1 confocal microscope, or a Zeiss LSM900 with AiryScan2 module.

#### Split-GFP immunohistochemistry

3-7-day old male fly brains were dissected in room temperature PBS and fixed in 4% paraformaldehyde solution at room temperature for 20 min. Fixed brains were then washed in PBST (0.3%) 3 x 20 min and blocked with normal goat serum (5%) for 30 min. The brains were then incubated with anti-Brp (nc82, 1:50, DSHB) with 5% goat serum for 2 d at 4 °C. No anti-GFP antibody was used. After 3 x 20 min washes in PBST, the brains were incubated with Alexa Fluor 546 goat anti-mouse (1:2000, ThermoFisher A11018) for 2 d at 4 °C. After 4 x 20 min washes in PBST, brains were mounted in Vectashield on a glass slide before scanning with a Nikon A1 confocal microscope.

Reconstituted split-GFP signal was quantified using ImageJ. The GFP signal was taken as the average pixel intensity within manually drawn volumes (freehand regions of interest in multiple z-slices) around the LC16 axon terminals and cell bodies. The background fluorescence (from an ROI in a proximal brain region outside of the LC16 neuron) was subtracted from the GFP signal. Statistical significance was evaluated by a t-tests and 2-way ANOVA in GraphPad Prism 9.

#### Connectomics search

We used the neuprint (hemibrain V.1.2.1 dataset) platform to search for candidate neurons and subsequent connectivity (https://neuprint.janelia.org/)

- Predicted link between LC16 and pC1a : Query Selection > General > Shortest paths > neuron A = LC16 # 1256830582 > Neuron B = pC1a # 359744514, Minimum weight =3
- 3D visualisation of 5-HT^PMPD01^ and pC1 neurons: “dataset”:“hemibrain:v1.2.1”,“bodies”[”297230760”,“\n297908801”,“\n359744514”,“\ n5813046951”,“\n267214250”,“\n267214250”,“\n392821837”,“\n359744514”,“\n581 3046951”,“\n514850616”]
- 3D visualisation of LC16 neurons and PPM1/2 neurons: “dataset”:“hemibrain:v1.2.1”,“bodies”[”1350945956”,“1288897930”,“1319927345”,“ 1319587380”,“1319579391”,“1254037524”,“1288893503”,“1289238972”,“13195868 61”,“1319919918”,“1412989088”,“950229431”,“792040520”,“5813054384”]

#### Statistics and Reproducibility

See Extended Data Table 1 and 2 for details on statistics. All statistical tests were performed using R or GraphPad Prism 9. Behavioural indexes and calcium imaging quantification are displayed as boxplots. Boxes delimit the lower (25th) and upper interquartile (75th) respectively, and the horizontal line represents the median. Each dot on the plot represents a single fly. Courtship progression behavioural data and locomotion data do not follow a normal distribution, thus nonparametric Mann-Whitney or Kruskal-Wallis test, followed by Conover- Iman multiple pairwise comparisons post-hoc test have been applied on raw data (p = 0.05, with a Bonferroni correction) for one factor experiments. In order to test the interaction between the genetic manipulations and the treatments we applied two factor ANOVA. Significant differences are indicated by different letters at the level of p<0.05. We employed a one-sample Wilcoxon signed rank test (μ=0) to assess if the speed change (Δ) in Extended Data Fig. 5e significantly deviated from 0. We indicated significance using * at the level of p<0.05.

Calcium imaging traces over time are represented as mean of ΔF/F0 (solid lines) with SEM (shaded area). Quantification plots are shown as min/max plots and median as horizontal line.

After verification of normality, paired t-test or paired Wilcoxon signed-rank test was applied on mean ΔF/F0 data from individual flies on specific time windows indicated in the graph and/or method sections. Significant differences are indicated by different letters (p < 0.05). Experimenters were not blinded to the conditions of the experiments during data collection. Genotypes used for one experiment were tested simultaneously and in random order as well as random times during the day to avoid any influence of circadian timepoints and order of the experimental trials. We repeated all statistical tests excluding data points that were identified as outliers using the ROUT method in PRISM with Q=0.5%, and always obtained the same results, so we did not exclude outlier data points.

## Data availability

Source data are provided with this paper.

## Code availability

Codes are available at https://github.com/lczl64/Cazale-Debat-Scheunemann-et-al..git

## Acknowledgements

We thank Johannes Felsenberg and Moshe Parnas for insightful comments and critical assessment of the manuscript. We thank Inbal Goshen, Tobias Hauser, Nuria Romero, Marta Moita and her lab for helpful discussions.

We thank Barry Dickson, Gerald Rubin, Mark Wu, Edward Kravitz, Julie Simpson, Andre Fiala, Anne Von Philipsborn, David Stern, Meghan Laturney, Suewei Lin, Yi Rao, Bowen Deng, Simon Sprecher, Hiromu Tanimoto, Jan Pielage, and Dana Galili for sharing fly stocks with us.

We would also like to thank Eve McCallion, Orla Cronin, Jasmine Stanley-Ahmed, Milan Narzary and Amber Kewin for help with fly and data collection and the Charité AMBIO Imaging Facility for support with live imaging. We also thank Julia Thüringer, Maria Luisa Vasconcelos and Saloni Rose for valuable discussions and help with data analysis. Finally, we thank members of the Rezaval and Scheunemann labs for useful comments on the manuscript.

This work was supported by Marie-Curie Sklodowska IF-EF-ST (101023536) to L.C.D; by DFG, under Germanýs Excellence Strategy – EXC-2049 – 390688087 to L.S. and D.O., 495407463 to L.S, 282979116 to D.O. and TP B04 of TRR265 (402170461) to D.O.; ERC (101088502) to D.O.; Wellcome Trust (225814/Z/22/Z), BBSRC (BB/S016031/1, BB/X000273/1, BB/X014568/1) to A.C.L; BBSRC (BB/W016249/1, BB/S009299/1), and Leverhulme Trust (RPG-2023-009) to C.R.

## Author information

These authors contributed equally: Laurie Cazalé-Debat, Lisa Scheunemann

## Authors and Affiliations

1School of Biosciences, University of Birmingham, Birmingham B15 2TT, UK

Laurie Cazalé-Debat, Megan Day, Anna Dimtsi, Youchong Zang, Lauren A Blackburn and Carolina Rezaval

2Freie Universität Berlin, Institute of Biology, Takustraße 6, 14195 Berlin

Lisa Scheunemann

3Charitè - Universitätsmedizin Berlin, Institut für Neurophysiologie and NeuroCure Cluster of Excellence, Chariteplatz 1, 10117 Berlin

Lisa Scheunemann, Tania Fernandez-d.V. Alquicira, Eric Reynolds and David Owald

4School of Biosciences, University of Sheffield, Firth Court, Western Bank, Sheffield S10 2TN, UK

5Neuroscience Institute, University of Sheffield, Firth Court, Western Bank, Sheffield S10 2TN, UK.

Katie Greenin-Whitehead, Andrew C. Lin

Present address:

6Biosciences Institute, Newcastle University, Newcastle upon Tyne, NE2 4HH, UK. 7Centre for Neural Circuits and Behaviour, University of Oxford, Oxford OX1 3SR, UK. 8School of Science and the Environment, University of Worcester, Worcester, WR2 6AJ. 9These authors contributed equally.

10These authors contributed equally.

*Corresponding author.

Contributions:

Conceived of and designed the study: L.C.D, and C.R; Methodology: L.C.D, L.S, D.O, A.C.L, and C.R.; Software: C.B; Investigation: L.C.D, L.S,M.D, T.F.V.A, A.D, Y.Z, L.A.B, K.G.W,

E.R, A.C.L. and C.R.; Resources: L.C.D, D.O, L.S and C.R.; Writing L.C.D, L.S., D.O, A.C.L. and C.R.; Supervision, L.C.D, A.C.L, D.O and C.R.

Corresponding author:

Correspondence to Carolina Rezaval

## Ethics declarations

*Competing interests*:

The authors declare no competing interests.

## TABLES

Extended Data Table 1: Statistics for behavioral, live calcium imaging and anatomical data (main and extended figures).

Extended Data Table 2: live calcium imaging inter-group comparisons

Extended Data Table 3: list of strains and genotypes

Extended Data Table 4: list of the optogenetic conditions per genotype and figure.

## Extended Data Figures

Extended Data Figure 1:

(A) Locomotion traces of solitary CS males before and after delivery of the threat (n=8).

(B) Change in average walking speed of 5–7-day old CS males before and after visual threat delivery (n=40).

(C) Percentage of males leaving the female (C1) and freezing index (C2) of males CS in absence (grey bar, n=59) or presence of a threat (blue bars, n=59) following 7 s of sustained courtship.

(D) Defensive index of males with TNT silenced LC16 neurons and genetic controls in absence (grey bars, n=18-38) or presence of the threat (blue bars, n=16-37).

(E) Defensive index after artificial activation of LC16 neurons under 660 nm light using CsChrimson in the absence of the threat (n^red^ ^light^ ^OFF^=15-16; n^red^ ^light^ ^ON^=20-27) and their genetic and light controls.

(F) Change in average walking speed of males with TNT silenced LC16 neurons and genetic controls before and after visual threat delivery (n=23-28).

(G) Courtship indexes of wild type and transgenic males exposed to a mechanical threat induced by a speaker emitting a loud 3 Hz binaural beat, causing surface vibrations (n=13-28).

(H) Left: Fluorescence intensity trace (ΔF/F0) over time of male flies expressing GCaMP7b in LC16 neurons pre and post exposure to a female fly under the microscope. Right: Quantification of mean ΔF/F0 comparing pre and post windows as indicated by pink rectangles below the traces (n=5).

Extended Data Figure 2:

**(A–B)** Courtship and defensive indexes of males with TNT silenced TRH neurons and genetic controls in absence (grey plots, n=24-33) or presence of the threat (blue plots, n=25-32).

**(C)** Change in average walking speed of males with TNT silenced TRH neurons and genetic controls before and after visual threat delivery (n=28-32).

**(D-E)** Courtship and defensive indexes of males with the TRH enzyme knocked out specifically in TRH neurons using RNA interference, and genetic controls in absence (grey plots, n=17-26) or presence of a late threat (blue plots, n=14-26).

**(F)** Change in average walking speed of males with the TRH enzyme knocked out specifically in TRH neurons and genetic controls before and after visual threat delivery (n=27-28).

**(G–H)** Courtship and defensive indexes after artificial activation of TRH neurons under 660 nm light using CsChrimson in the absence of the visual threat (n^red^ ^light^ ^OFF^=25-32; n^red^ ^light^ ^ON^=25-31) and their genetic and light controls.

Extended Data Figure 3:

**(A)** Courtship index of P1 neurons expressing CsChrimson without the 660 nm red light, in absence (grey bars, n=25-31) or presence (blue bars, n=31-35) of the visual threat.

**(B)** Fluorescence intensity trace (ΔF/F0) over time of males expressing GCaMP6m in TRH^PMPD^ neurons under 1040 nm two-photon laser stimulation. Lower Left: Quantification of mean ΔF/F0 of baseline activity compared to stimulation window (2 s time window each) (n=9). Lower right: image of the focal plane showing region of interest during imaging (scale bar: 15µm) referring also to Fig. 2e.

**(C)** Change in average walking speed of TNT silenced TRH^R23E12^ neurons and genetic controls before and after visual threat delivery (n=31-40).

**(D)** Defensive index of males with TNT silenced TRH^R23E12^ neurons and genetic controls in absence (grey bars, n=15-17) or presence of the threat (blue bars, n=14- 24).

**(E–F)** Courtship and defensive indexes of males with the TRH enzyme knocked down specifically in TRH^R23E12^ neurons using small RNA interference, and genetic controls in absence (grey plots, n=18-25) or presence of a late threat (blue plots, n=18-25).

**(G)** Defensive index after artificial activation of TRH^R23E12^ neurons under 660 nm light using CsChrimson in the absence of the visual threat (n^red^ ^light^ ^OFF^=17-25; n^red^ ^light^ ^ON^=14- 22) and their genetic and light controls.

**(H)** Behavioral effects of knocking down 5-HT receptors in P1 neurons via small RNA interference, in control (n=38) and experimental males (n=18-27).

**(I)** Behavioral effects of knocking down Gɑi and Gɑo proteins in P1 neurons via small RNA interference (n=17), and controls (n=12).

**(J)** Upper panel: Fluorescence intensity trace (ΔF/F0) over time of male flies expressing GCaMP6s in P1 neurons pre and post application of 100 µM 5-HT and quantification of mean ΔF/F0 comparing the 30 s pre and post 30 s post time windows (n=6). Lower panel: Fluorescence intensity trace (ΔF/F0) over time of male flies expressing GCaMP6s in P1 neurons and simultaneous knock down of Gɑi protein pre and post application of 100 µM 5-HT. Quantification of mean ΔF/F0 comparing the 30 s pre and post 30 s post time windows (n=6).

**(K)** Change in average walking speed of males with either the 5-HT7 serotonin receptor or the Gɑi protein knocked out specifically in P1 neurons before and after visual threat delivery (n=31-40).

Extended Data Figure 4:

**(A–B)** Courtship (A) and defensive (B) indexes without red light (left panel) or after artificial activation of plP10 descending neurons under 660 nm red light (right panel) using CsChrimson in absence (grey bars, n=30) or presence (blue bars, n=28-30) of the visual threat.

**(C–D)** Courtship (C) and defensive (D) indexes after artificial inhibition of plP10 descending neurons under 515nm light using the green-light activatable anion channelrhodopsin GtACR1 in the absence of the visual threat (n^green^ ^light^ ^OFF^=22-25; n^green^ ^light ON^=20-24) and their genetic and light controls.

**(E)** 5-HT RNAi receptor screen: courtship indexes of plP10 Split-GAL4 controls (n=35) and males with one of the five subtypes of the *Drosophila* 5-HT receptor knocked down specifically in P1 neurons using small RNA interference (n=7-36).

**(F)** Change in average walking speed of males the 5-HT2B serotonin receptor knocked out specifically in plP10 neurons and controls before and after visual threat delivery (n=18-20).

**(G)** Behavioral effects of knocking down Gɑi and Gɑo proteins in plP10 neurons via RNAi and controls (n=12-18).

**(H)** Upper panel: Fluorescence intensity trace (ΔF/F0) over time of male flies expressing GCaMP6s in plP10 neurons pre and post application of 100 µM 5-HT and quantification of mean ΔF/F0 comparing the 30 s pre and last 30 s post time windows (n=5). Lower panel: Fluorescence intensity trace (ΔF/F0) over time of male flies expressing GCaMP6s in plP10 neurons and simultaneous knock down of Gɑo protein pre and post application of 100 µM 5- HT. Quantification of mean ΔF/F0 comparing the 30 s pre and last 30 s post time windows (n=7).

Extended Data Figure 5:

**(A-B)** Fraction of flies exhibiting courtship or defensive behavior in response to an early

(A) or late (B) threat, delivered after 7 s or 4 min of sustained courtship (n=45).

**(C)** Behavioral protocol: the threat is delivered at either 30 s, 5, 10 or 15 min after copulation initiation. Controls for copulation have been observed at the same time point in absence of the threat. (Note that sperm transfer takes ∼10 min: Crickmore and Vosshall, 2013.)

**(D)** Percentage of flies engaged in copulation 30 s, 5, 10 or 15 min after mating initiation in absence or presence of the threat. The threat is delivered for 30 s at the aforementioned time points.

Extended Data Figure 6:

(A) Courtship index of TH-C1 GAL4 neurons expressing CsChrimson without the 660 nm red light, in absence (grey bars, n=15-21) or presence (blue bars, n=18-23) of the visual threat.

(B) Defensive index without red light (left panel) or after artificial activation of TH-C1 neurons under 660 nm red light (right panel) using CsChrimson in absence (grey bars, n=15-21) or presence (blue bars, n=17-23) of the visual threat.

(C) Difference in walking speed before and after the threat of solitary males and solitary females with TH-C1 neurons artificially activated under 660 nm red light during the threat delivery using CsChrimson (purple plots, n=28-52) and its light controls (grey plots, n=54-64).

(D) Ethograms of abdomen bending events of male TH-C1>GCaMP7b flies during the 4 min period exposed to a female under the microscope.

(E) Quantification of abdomen bending events comparing the first tier (0–80 s), second tier (80-160 s) and third tier of the 4 min period of male flies courting a female under the microscope (n=10).

(F) Fluorescence intensity trace (ΔF/F0) over 4 min in male flies expressing GCaMP7b in PPM1/2 neurons while paired with a male fly under the microscope. Middle: Quantification of mean ΔF/F0 at 1 min compared to 4 min (20 s time window each) (n=6). Right: Comparison of full abdomen bending events when males were paired with females or males under the two-photon microscope (n=10).

(G) Fluorescence intensity trace (ΔF/F0, blue) over 7 min of males expressing GCaMP7b in PPM1/2 dopamine neurons while courting a female under the microscope and control ROI trace of an adjacent neuron from the same acquisition (grey). The female was approached after 30 s and removed after 4 min. Right panel: Quantification of mean ΔF/F0 comparing intensities at 1 min, 4 min and 7 min time windows (10 s each) (n=6).

(H) Fluorescence intensity trace (ΔF/F0) over 4 min of males expressing GCaMP7b in PAL neurons while courting a female under the microscope. Right panel: Quantification of mean ΔF/F0 comparing 1 min to 4 min time windows (20 s each) (n=6). Images below show representative fluorescence heatmap of PAL neurons at timepoints 1 min, 2 min, 3 min and 4 min (scale bar 6µm).

(I) Fluorescence intensity trace (ΔF/F0) over 4 min of males expressing GCaMP7b in PAM neurons while courting a female under the microscope. Right panel: Quantification of mean ΔF/F0 comparing 1 min to 4 min time windows (20 s each) (n=6). Images below show representative fluorescence heatmap of PAM neurons at timepoints 1, 2,3 and 4 min (scale bar 6µm).

(J) Alignment of abdominal bending events during neural recording of PPM1/2 neurons in a TH-C1>GCaMP7b male paired with a virgin female.

(K) Fluorescence intensity trace (ΔF/F0) over 4 min in male flies expressing GCaMP7b in PPM1/2 with a fixed unmovable proboscis while paired with a female fly under the microscope. Right: Quantification of mean ΔF/F0 at 1 min compared to 4 min (20 s time windows each) (n=5).

(L) Fluorescence intensity trace (ΔF/F0) over 4 min in male flies expressing GCaMP7b in PPM1/2 with a fixed unmovable front legs while paired with a female fly under the microscope. Right: Quantification of mean ΔF/F0 at 1 min compared to 4 min (20 s time windows each) (n=5).

(M) 660 nm red LED stimulation protocol of Fig.4f1-3 and number of evoked abdominal bendings of male THC1>GCaMP6s; OvAbg>CsChrimson flies within the pre-stimulus, during-stimulus and post-stimulus windows (n=5-6).

(N) Number of courtship events (tapping or licking) of males THC1>GCaMP7b paired with a female during PPM1/2 neural recording under the two-photon microscope comparing control males with males unable to move the abdomen (n=10-6).

Extended Data Figure 7:

(A) Fluorescence intensity trace (ΔF/F0) over 4 min in male flies expressing the GRABDA sensor in LC16 neurons when paired with a male fly under the microscope. Right: Quantification of mean fluorescence intensity at 1 min compared to 4 min (20 s time window each) (n=5).

(B) Fluorescence intensity trace (ΔF/F0) over time in male flies expressing GCaMP6f in LC16 neurons pre and post application of 500 µM dopamine. Right: Quantification of mean fluorescence intensity comparing the 30 s pre and last 30 s post time windows (n=10).

(C) Fluorescence intensity trace (ΔF/F0) over time of male flies expressing GCaMP7b in LC16 neurons following focal injection of either dopamine (blue) or saline (grey) to LC16 axons or dopamine in flies co-expressing Dop2R-RNAi (purple). Arrow shows time point of injection. Example images of focal dopamine injection; white dashed outline shows region of interest (LC16 axons); injection pipette (with Texas Red) is labelled in magenta. Scale bar: 5 µm. Right: Quantification of mean ΔF/F0 comparing the pre and post injections time windows. Saline (n=5 hemispheres from 3 flies) v. Dopamine (n=6 hemispheres from 4 flies, Dop2R-RNAi (n=5 hemispheres from 3 flies).

(D) TH genomic region indicating sequences upstream of Gal4 for lines C1 and C’. Exons are shown as black boxes. Translational start, stop, and introns sites are named ATG, stop and i1- i6, respectively. The “+”, “+/-”, and “-” in the table report the expression of C1 and C’ in most, a subset, or none of the neurons in the clusters, respectively. Schematic from ^108^ S2. **(E1-3)** Fluorescence intensity trace (ΔF/F0) over time of male flies expressing LexAop GCaMP6f in LC16 neurons and UAS-CsChrimson in PPM1/2 neurons pre and post either 30 s of: **E1** threat exposure under the microscope, **E2**: threat exposure in presence of 15x LED stimulations for 1 s, or **E3**: threat and LED stimulation in control flies expressing only UAS- GCaMP6f in LC16 neurons. Right: Quantification of mean ΔF/F0 comparing the 15 s pre and 30 s post time windows (n=7-8). Note that the control’s threat response in LC16>GCaMP6f; PPM1/2>CsChrimson flies without the LED is comparatively low. Thus, we might not be able to detect graded modulation of LC16 neurons by activation of PPM1/2 neurons.

(F) Above: UAS-spGFP1-10 expressed under the control of LC16 Split-GAL4 without (left) or with (right) tagging endogenous Dop2R with spGFP11 (Dop2R::GFP11). Below: quantified GFP fluorescence in the LC16 cell bodies (left) or axon terminals (right) where UAS-spGFP1- 10 is expressed under the control of LC16 Split-GAL4, with or without Dop2R::GFP11. Neuropil counterstained with anti-brp (nc82; magenta). Dashed white outlines represent the regions of interest for the axon terminals and cell bodies of the LC16 neurons (n=8-10 brains). GFP fluorescence was normalised to the average fluorescence in the LC16 axon terminals in LC16>GFP1-10,Dop2R::GFP11 flies. Scale bar: 25 μm. Right panels are reproduced from main Fig. 5i for comparison.

(G) Fluorescence intensity trace (ΔF/F0) over time of male flies expressing GCaMP6f in 5- HT^PMPD^ neurons pre and post (1) 30 s of threat exposure under the microscope; (2) threat exposure in presence of 500 µM dopamine and (3) threat exposure in presence of 500 µM dopamine with additional knock down of Dop2R in LC16 neurons. Right: Quantification of mean ΔF/F0 comparing the 15 s pre and 30 s post time windows (n=5-7).

## Supplementary Videos

Supplementary Video 1: Wild-type CS male displaying early courtship steps toward an immobile female.

Supplementary Video 2: Wild-type CS male displaying late courtship steps toward an immobile female.

Supplementary Video 3: Wild-type CS male displaying freezing behavior in response to an early threat, delivered 7 s after courtship initiation.

Supplementary Video 4: Wild-type CS male running away in response to an early threat, delivered 7 s after courtship initiation.

Supplementary Video 5: Wild-type CS male pursuing courtship and displaying abdominal bending in the presence of a late threat, delivered 4 min after courtship initiation.

Supplementary Video 6: Tethered wild-type male CS displaying abdominal bending toward a female under the two-photon microscope.

Supplementary Video 7: Tethered TH-C1> GCaMP7bmale with fixed abdomen paired with a female under the two-photon microscope.

Supplementary Video 8-9: Tethered TH-C1>GCaMP7b males with either a fixed proboscis or fixed front legs, respectively, paired with a female under the two-photon microscope.

## Notes

### Competing Interest Statement

The authors have declared no competing interest.

