## Supplementary figures and images for "Mating proximity blinds threat perception"

### EXTENDED DATA FIGURE 1

#### Extended Data Figure 1

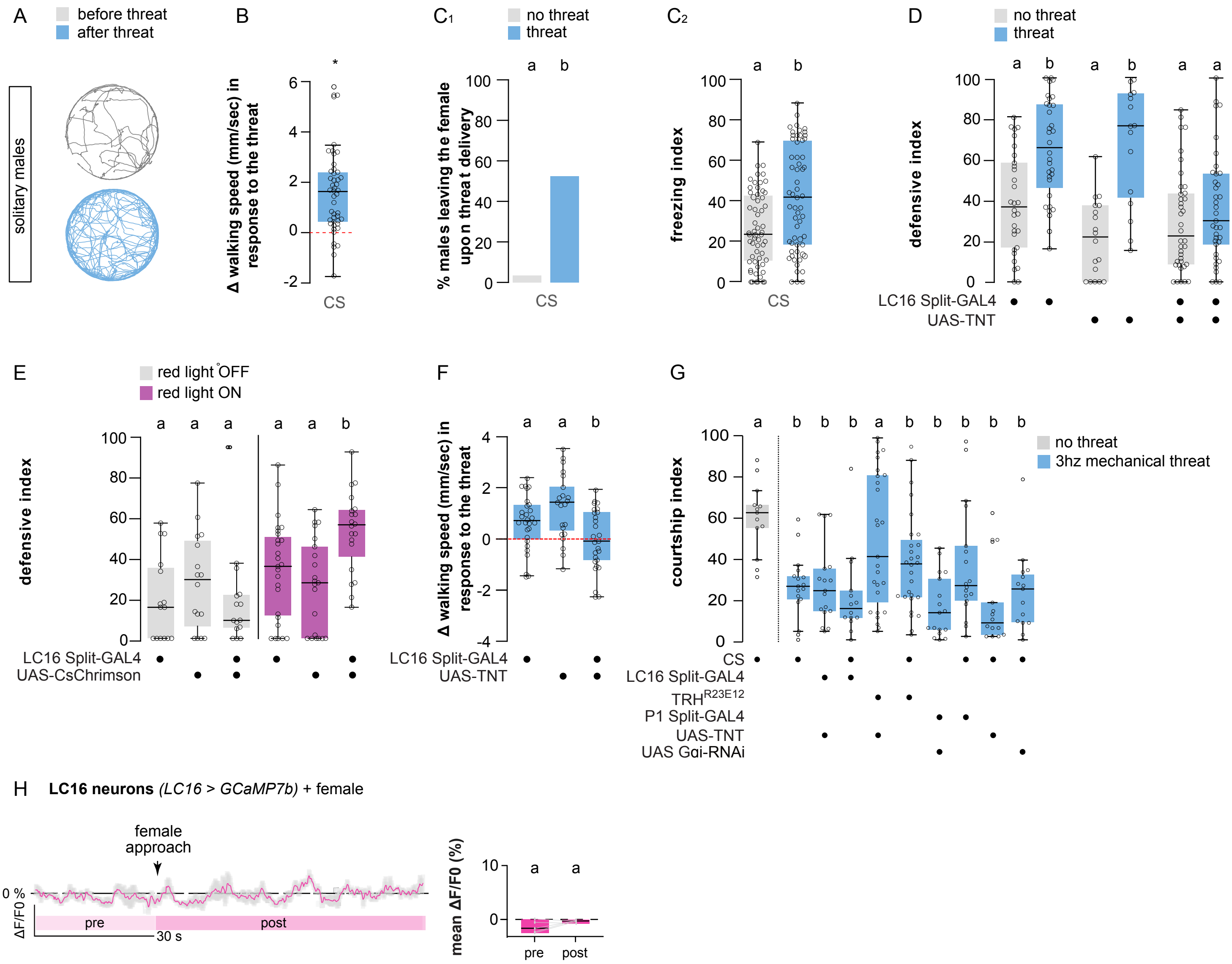

### EXTENDED DATA FIGURE 2

#### Extended Data Figure 2

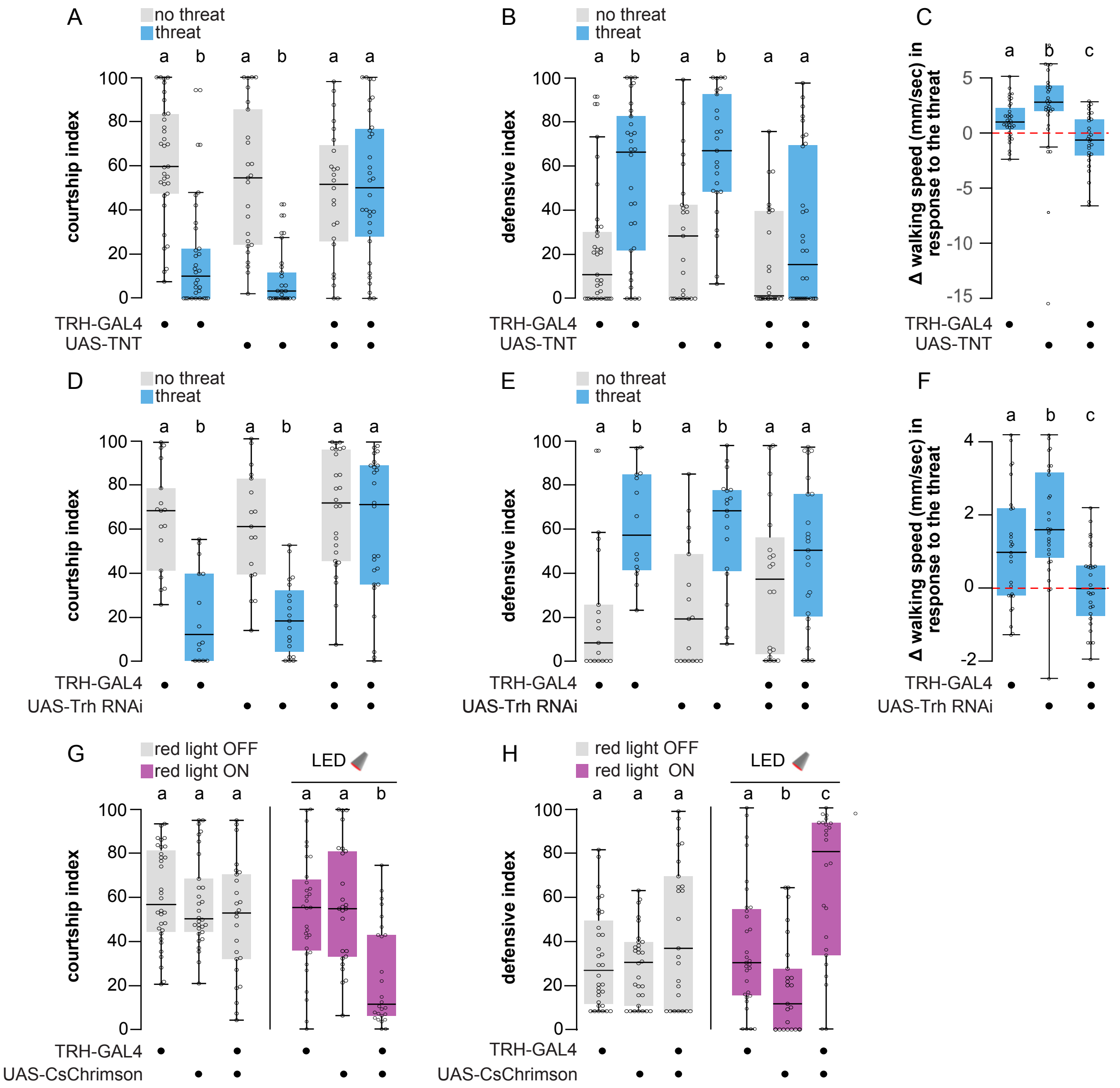

### EXTENDED DATA FIGURE 3

#### Extended Data Figure 3

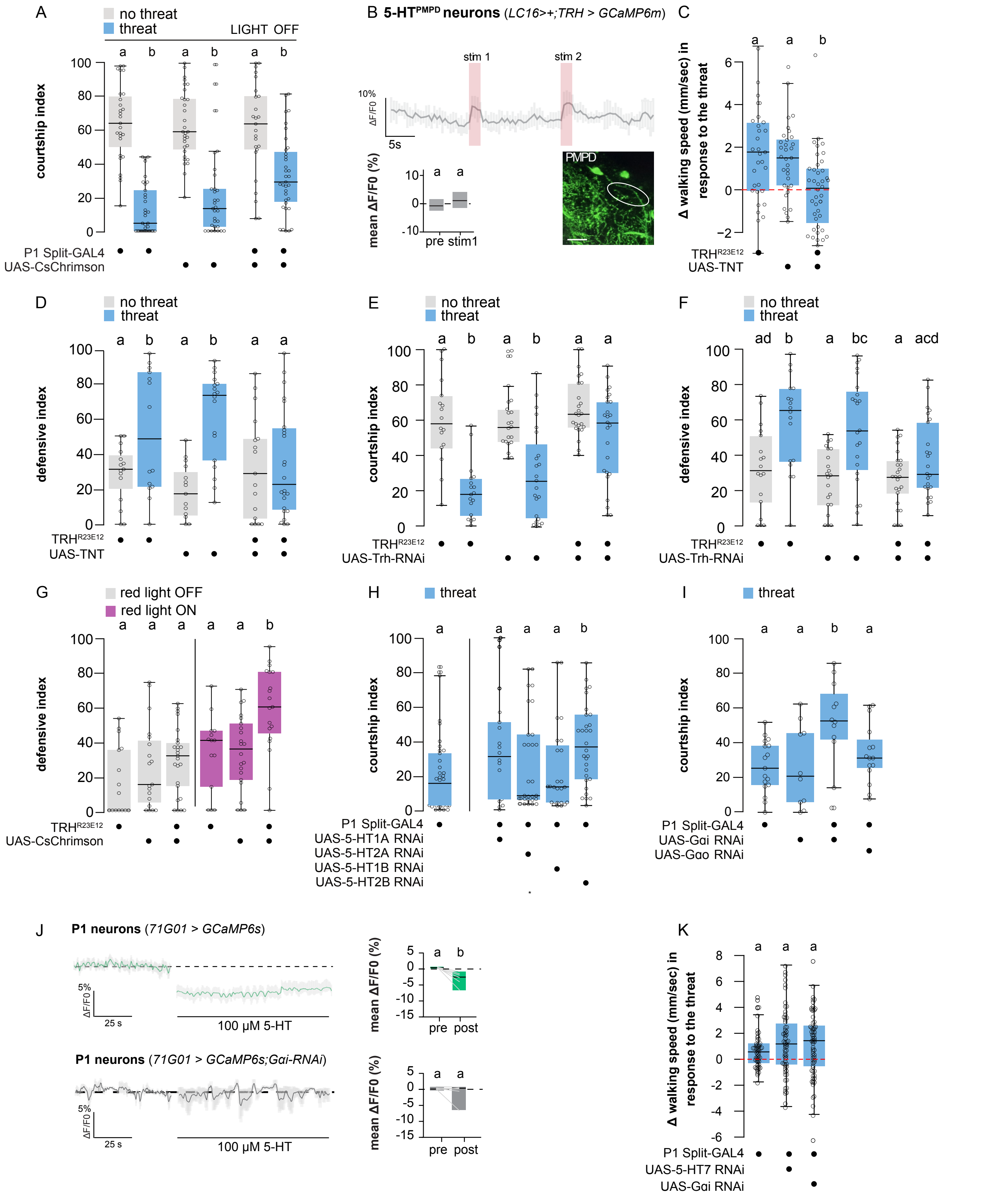

### EXTENDED DATA FIGURE 4

#### Extended Data Figure 4

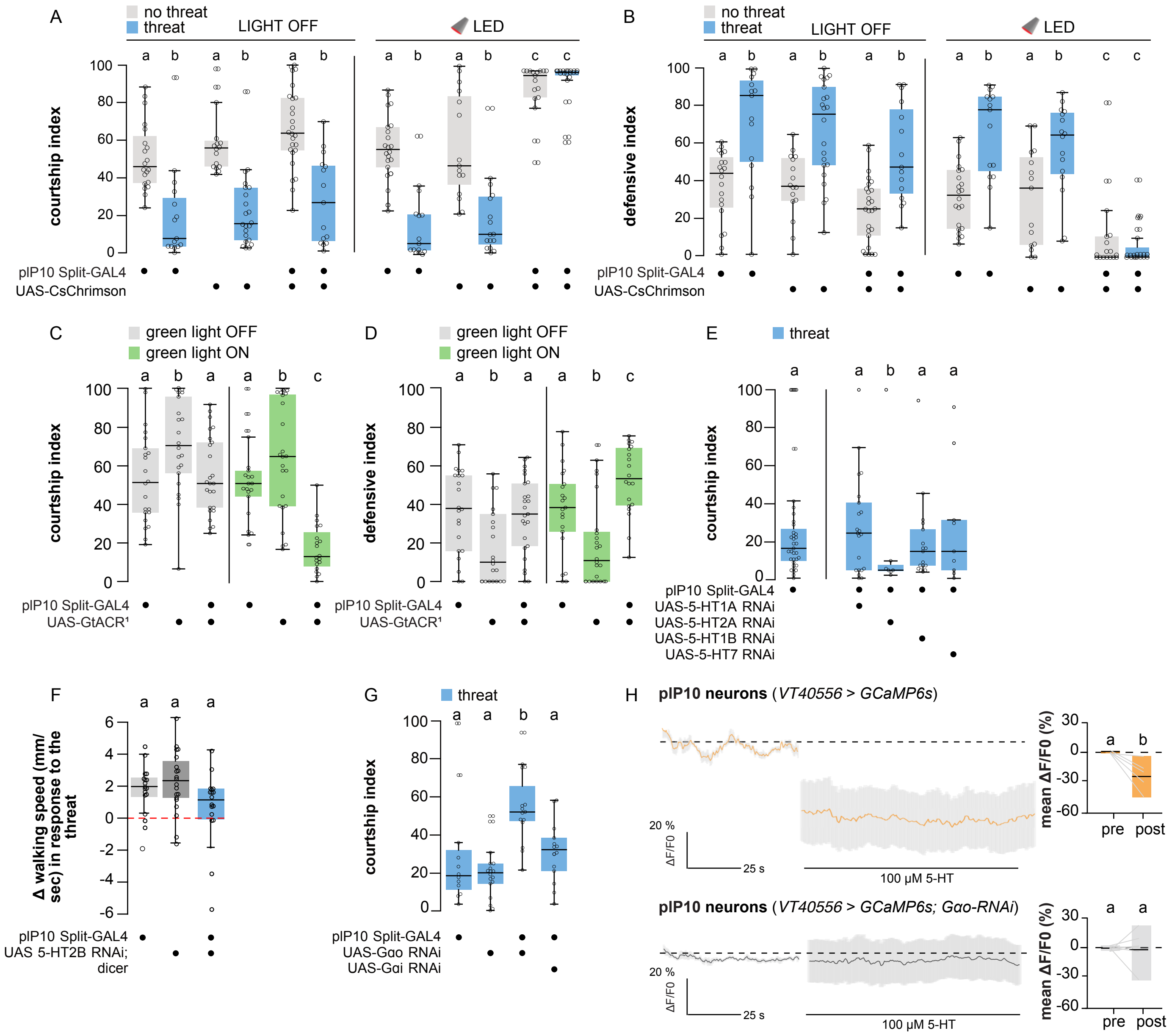

### EXTENDED DATA FIGURE 5

#### Extended Data Figure 5

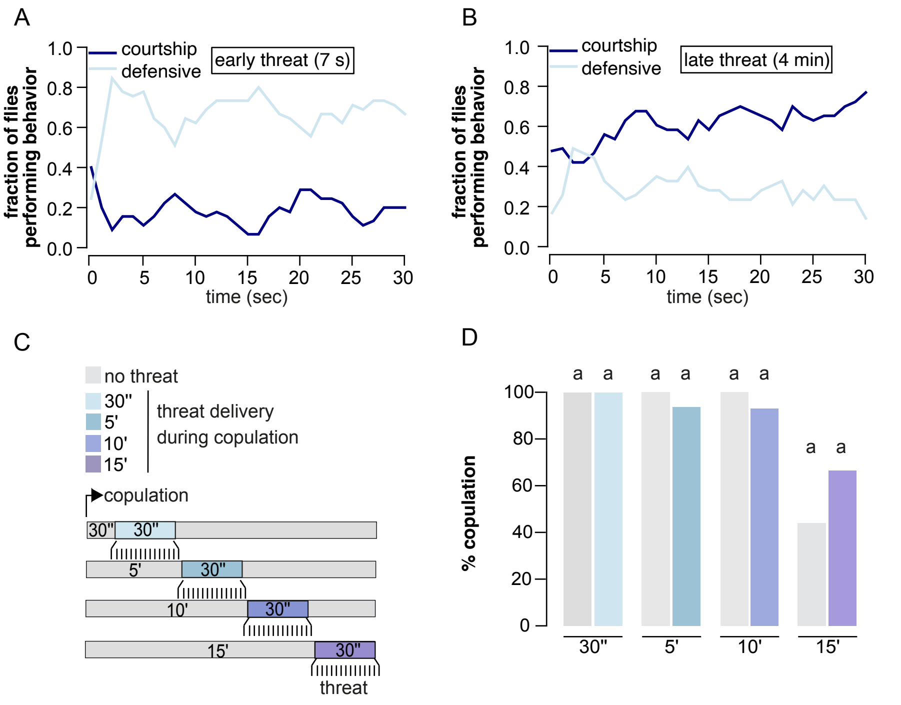

### EXTENDED DATA FIGURE 6

#### Extended Data Figure 6

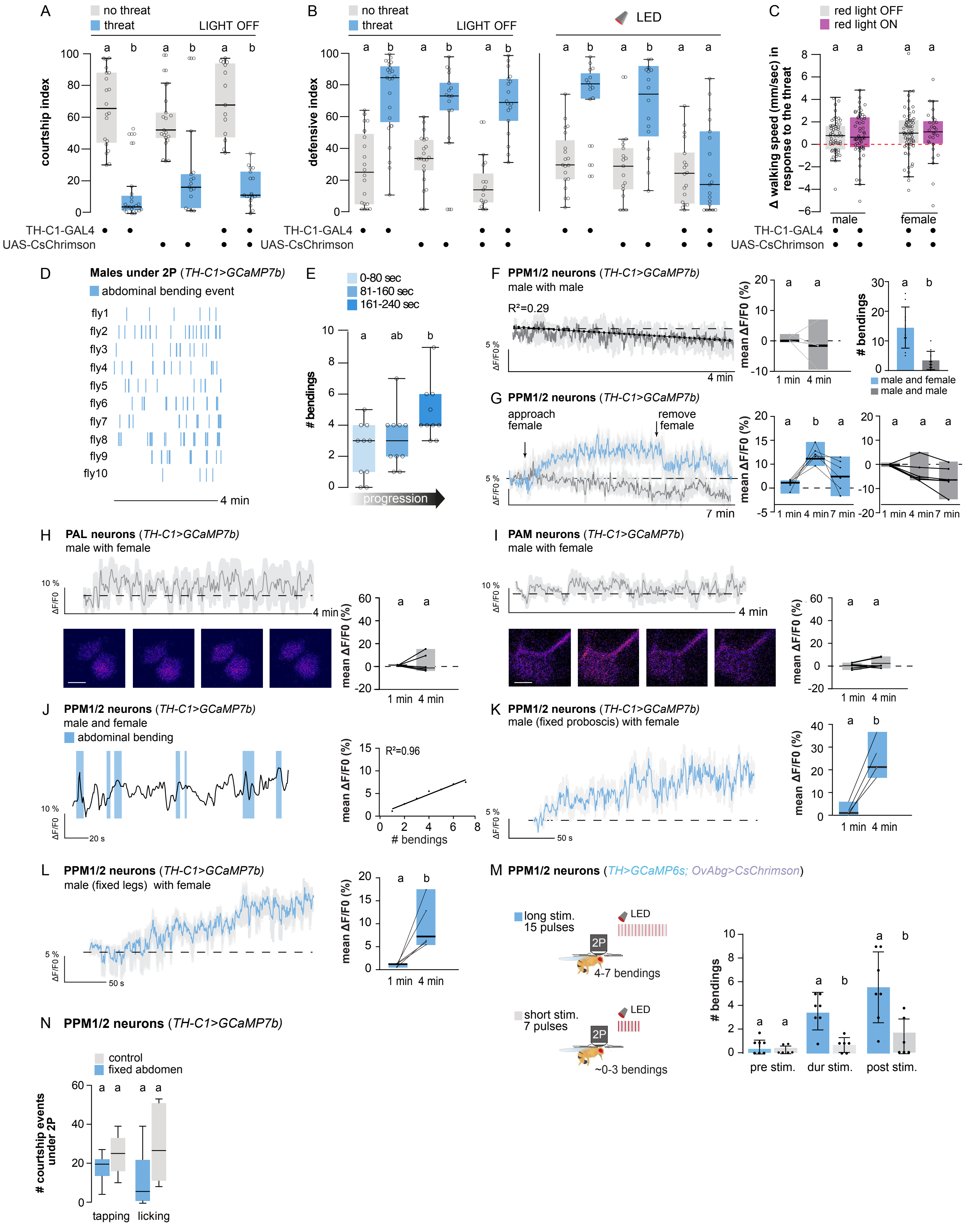

### EXTENDED DATA FIGURE 7

#### Extended Figure 7

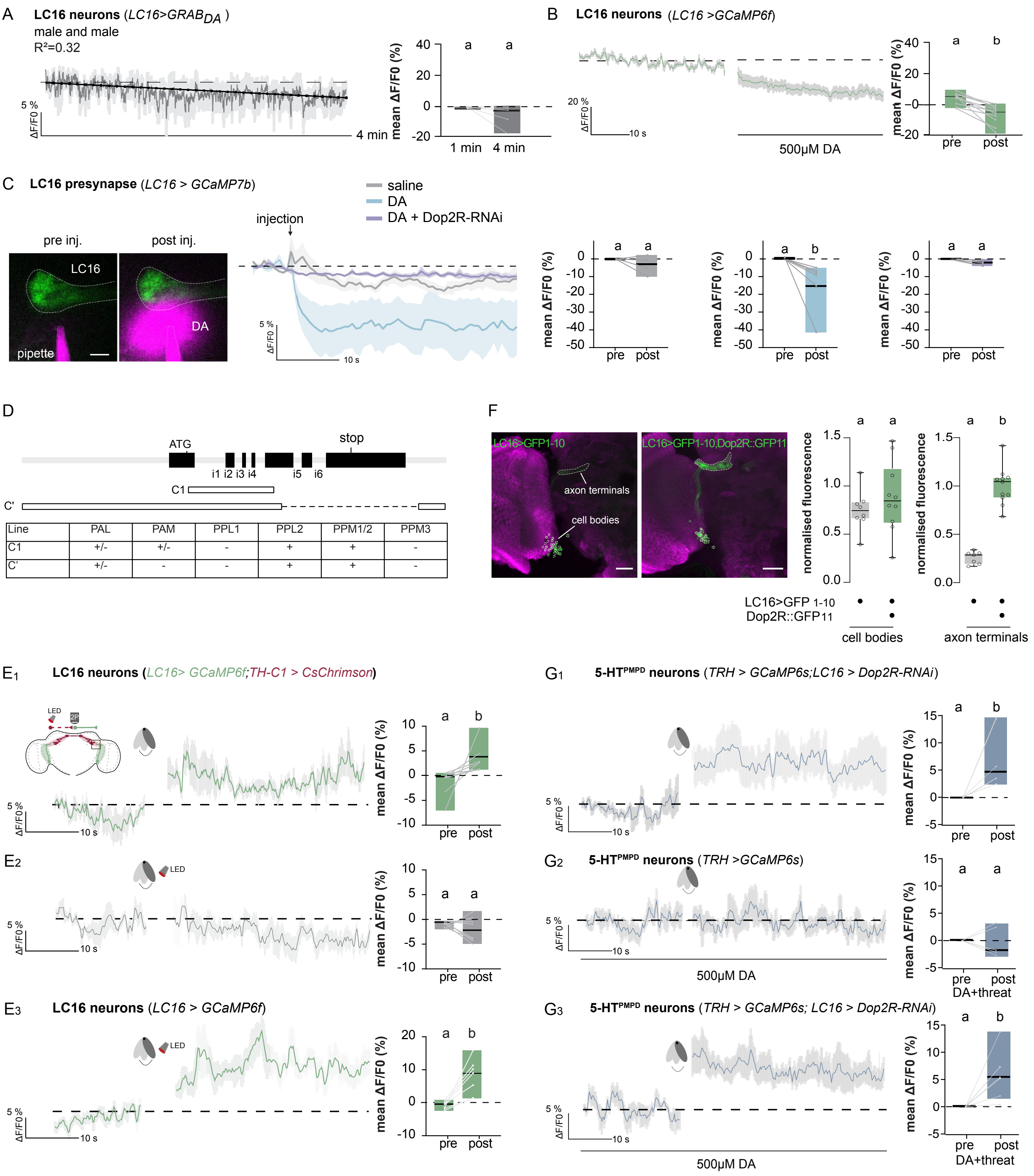
